# Plasticity Associated with Adoption of Social Roles in Clown Anemonefish

**DOI:** 10.64898/2025.12.09.693150

**Authors:** Lili F. Vizer, Douglas Alvarado, Colleen Bove, Marcela Herrera, Annabel Hughes, Kian Thompson, Steven M. Bogdanowicz, Vincent Laudet, Sarah W. Davies, Peter M. Buston

**Affiliations:** Department of Biology, Boston University, 5 Cummington Mall, Boston, MA, USA 02215; Marine Eco-Evo-Devo Unit, Okinawa Institute of Science and Technology, 904-0495 Onna-son, Okinawa, Japan; Department of Ecology and Evolutionary Ecology, Cornell University, Ithaca, NY, USA 14853-2701

**Keywords:** phenotypic plasticity, growth, feeding behavior, agonistic behaviors, gene expression, anemonefish, Amphiprion

## Abstract

Central questions of animal behavior are how do individual phenotypes shape and how are individual phenotypes shaped by dominance hierarchies. Using clownfish (*Amphiprion percula*) as a model system, we investigated how individuals strategically modify their phenotype during hierarchy formation. We tested the overarching hypothesis that interactions among size- and age-matched rivals will lead to the emergence of social roles (dominant, subordinate, solitary), accompanied by divergence in four aspects of individual phenotypes: growth, feeding behavior, agonistic behaviors (aggression, submission), and gene expression. To test this, we created 30 replicates of juvenile clownfish, each comprising a size-matched pair housed together, and a solitary size-matched individual housed separately. Our results show that an individual’s social context shapes its growth trajectory through coordinated changes in feeding behavior, agonistic behaviors, and gene expression. Individuals emerging as dominant within pairs grew twice as much compared to those emerging as subordinates, while subordinate and solitary individuals showed comparable growth. Initially, paired individuals consumed more food per capita than solitary individuals, but this difference declined as size hierarchies became established. Agonistic behaviors also decreased over time among paired individuals as size differences emerged. Transcriptomic analyses revealed upregulation of conserved vertebrate growth pathways and ossification related genes, and downregulation of satiety-associated genes, in dominant individuals compared to subordinates and solitaries. Here we identify associations between changes in gene expression, growth, and feeding behavior regulation that reinforce social role differentiation during dominance hierarchy formation in clownfish. Our findings lay the foundation for a broader framework to explore the mechanisms underpinning strategic growth in social vertebrates.

## Introduction

Social groups, in which some individuals forgo their own reproduction and help others to reproduce, are found in a wide variety of taxa (Rubenstein & Abbot, 2017). Within such social groups, dominance hierarchies in which individuals adopt particular social positions, are ubiquitous (Chase, 1982). Dominance hierarchies are determined by and subsequently determine interactions among group members (Milewski et al., 2022). Such hierarchies are commonly based on dominance and submission behaviors, where certain individuals of higher social rank often hold power over and are aggressive toward submissive individuals in lower social ranks (Tibbetts et al., 2022). While dominance hierarchies are often associated with asymmetric aggression toward subordinates, the structure, the overall number of members, and the number of rivals at any given social rank can vary within and among species (Drews, 1993). Regardless of the exact nature of the hierarchy, dominant individuals typically enjoy priority access to limited resources and can monopolize food, mates, and/or shelter, while subordinate members may have to wait their turn or accept less desirable resources (Chase, 1982).

Being in a dominant position confers advantages leading to higher fitness, so selection is expected to favor traits that support the acquisition of dominant positions. Social ranks, and thus the acquisition of a dominant position, at the establishment of dominance hierarchy are predominantly determined by costly (in terms of risk of injury, time, and energy invested) antagonistic interactions (Chase et al., 2002; Tibbetts et al., 2022; Vieira & Peixoto, 2013). During these interactions, phenotypic differences, such as fighting capacity, morphology, and physiology, will largely determine winners and losers (Tibbetts et al., 2022). Winners tend to be larger in size (López-Segoviano et al., 2018; Thornton et al., 2015; Wong et al., 2016), be more aggressive (Hobson et al., 2021), while losers tend to be smaller in size (Buston, 2003; Johnston et al., 2021), and submissive (Hobson et al., 2021; Tibbetts et al., 2022). Winners and losers may also differ in their feeding behavior (Huchard et al., 2016; Wong et al., 2008). These observations have generated a traditional view of dominance hierarchies where the phenotypes of individuals determine their social position.

Once the initial conflict over social position is resolved, selection will favor individuals who adjust their phenotypes in a way that maximizes the benefits in their new social position and minimizes the costs of potential future conflicts. Indeed, empirical data suggest that most hierarchies are somewhat stable over time (Gronenberg et al., 2008; Tibbetts et al., 2022), and this stability can be attributed to phenotypic changes as members take on fixed social roles leading to distinctly different phenotypes across social ranks. These distinct phenotypes are adjusted as individuals ascend or descend within the hierarchy, thus social rank influences phenotypes of individuals. For example, in naked mole-rats (*Heterocephalus glaber)*, when the queen dies, non-breeding subordinates often engage in violent competition for her role. Individuals that acquire the dominant position will undergo rapid sexual maturation and growth, including elongation of the lumbar vertebrae, to become the new queen; other individuals keep their subordinate positions, and remain sexually immature and small (Johnston et al., 2021). This example highlights that adjustment of phenotypes to fit social roles is favored, playing a crucial role in the maintenance of dominance hierarchy stability. Nevertheless, processes where social rank drive the phenotype of members within a dominance hierarchy are less well understood and often overlooked, as it can be challenging to disentangle whether a phenotype of an individual is the cause or the consequence of their social rank without long-term investigations of marked individuals (Milewski et al., 2022).

Understanding the proximate mechanisms that facilitate phenotypic adjustments is key to disentangling whether phenotypes are the cause or consequence of social rank, yet relatively few studies have addressed this question. This is further complicated by the fact that, in social vertebrates, phenotype (including gene expression) can both influence and be influenced by social interactions, often through continuous feedback loops rather than a clear directional relationship. Research in social mammals, such as the naked mole rats (Dengler-Crish & Catania, 2007), and Damaraland mole rats (Johnston et al., 2021; Thorley et al., 2018) as well as in social fish (Maruska & Fernald, 2014; Renn et al., 2008), has shown that shifts in social rank are accompanied by complex changes in hormonal and neural pathways, which lead to the observed coordinated morphological shifts. In Damaraland and naked mole-rats, ascension to queen social role involves changes in behavior in conjunction with rapid skeletal remodeling that occurs without the need of pregnancy (Dengler-Crish & Catania, 2007). Hormonal profiling further revealed that estrogens and androgens play nuanced, context-dependent roles in relation to an individual’s social status (Johnston et al., 2021). In cichlids, transitions from subordinate to dominant males are characterized by immediate early gene activation in the brain, followed by changes in coloration, growth, and sexual maturation, alongside dynamic regulation of the stress axis (Alward et al., 2020; Culbert et al., 2018; Hofmann & Fernald, 2000; Maruska & Fernald, 2014; Renn et al., 2008). Despite these insights, our understanding of proximate mechanisms remains limited. Most studies, often conducted without relevant social context of an individual or in species entirely lacking dominance hierarchies, have examined single pathways to uncover variation in coloration (Salis et al., 2019, 2021), appetite regulation (see review for teleosts: Volkoff, 2019), behavior (Bender et al., 2006; Renn et al., 2008; Santema et al., 2013; Solomon-Lane et al., 2022; see review for teleosts: St-Cyr & Aubin-Horth, 2009), and growth (Beckman, 2011; Lu et al., 2020; see reviews for teleosts: Reinecke, 2010; Zhou et al., 2024), leaving the broader molecular shifts associated with social role adoption unresolved. Gene expression profiling offers a powerful approach to uncover coordinated changes in gene expression across multiple pathways, providing insight into associations between the development of social role-specific phenotypes and their underlying proximate mechanisms.

The clown anemonefish, *Amphiprion percula*, provides an ideal system to investigate both how an individual’s phenotype influences its social rank and how its social rank influences its phenotype during the formation and maintenance of dominance hierarchies. Its well-annotated genome (Lehmann et al., 2019) enables transcriptomic analyses that can help characterize the molecular patterns associated with these processes. Clown anemonefish live in close association with sea anemones, with one social group per anemone. Each social group is composed of a breeding pair and 0-4 non-breeders. Within each group, there is a size-based dominance hierarchy: the female is largest (rank 1), the breeding male is second largest (rank 2), followed by non-breeders (rank 3-6) (Buston, 2003, 2004b; Buston & Cant, 2006). This size hierarchy represents a queue for breeding positions: when the female of the group dies, the breeding male changes sex and assumes the position vacated by the female, and in turn, the largest non-breeder ascends to the breeding male role by growing and becoming sexually mature (Buston, 2003, 2004a; Mitchell, 2005). Whenever a rank is vacated, the individual next in line ascends by growing and modifying its phenotype to fulfill the new social role, triggering a cascade where all lower-ranked individuals advance one rank up in a similar manner. Underlying this cascade is an underappreciated aspect of this system (and likely many other systems) where phenotype (including gene expression) influences the outcome of social interactions, and the outcome of social interactions influences the phenotype (including gene expression). While some aspects of clownfish social dynamics are well-understood, other phenotypic, physiological, and underlying changes in gene expression associated with the formation and maintenance of the social hierarchy of *A. percula* remain unexplored.

Here, our goal is to determine how phenotypes of size-matched and age-matched rivals change at the initiation of size-based dominance hierarchies in clownfish. In other words, how members adjust their phenotypes as they grow into their respective social roles as dominant and subordinates, and how their ultimate gene expression patterns reflect these phenotypes. Given that an individual’s social role is dependent on its social position (dominant, subordinate, solitary), we tested the overarching hypothesis that interactions among size- and age-matched rivals will lead to the emergence of social roles accompanied by divergence in four aspects of individual phenotypes: growth, feeding behavior, agonistic behaviors, and gene expression (Fig. 1). We predicted that paired size-matched rivals will grow faster than size-matched solitary individuals, and that one of the pair will grow faster than the other, as they attempt to outgrow each other (Desrochers et al., 2020; Huchard et al., 2016; Reed et al., 2019). In parallel, we predicted that paired size-matched rivals will eat more than solitary individuals as they attempt to outgrow each other (Huchard et al., 2016). Within size-matched rival pairs, we also predicted that agonistic behaviors (aggression, submission) will decline over time, as size differences emerge (Wong et al., 2016). Finally, we predicted that individuals taking on dominant roles will show upregulation of genes associated with ossification, growth, and appetite stimulation, consistent with their enhanced growth trajectories (Lu et al., 2020; Volkoff, 2019; Zhou et al., 2024).

**Figure 1:**
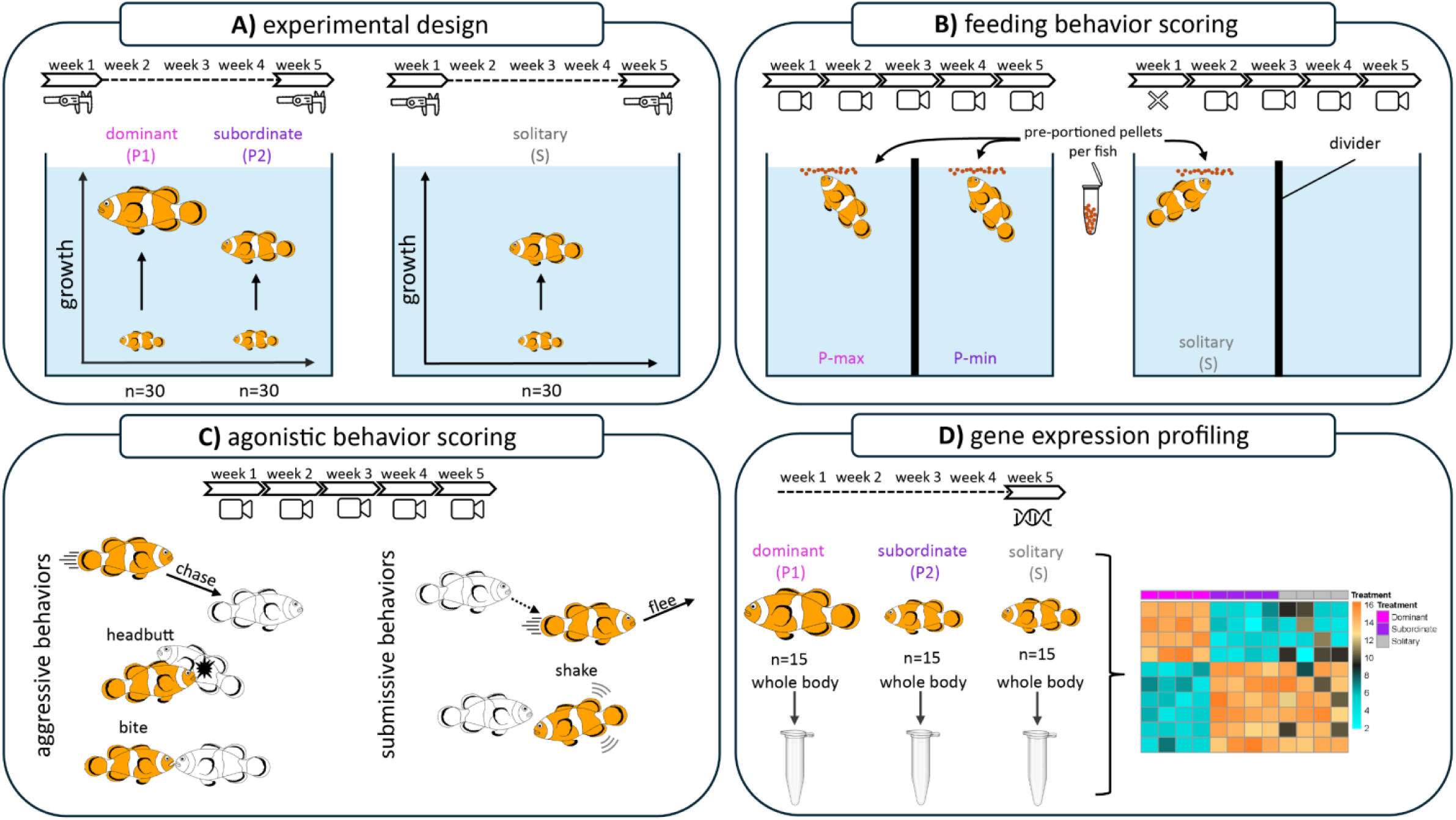
Overview of experimental design and methods. **A)** *Schematic of the experimental design.* Replicate groups of juvenile clownfish (*Amphiprion percula*) housed as size-matched pairs (n = 30 pairs) and size-matched solitary individuals (n = 30 individuals). Standard length (SL) was measured at the beginning and end of the five-week experiment, and individuals were denoted as dominant (P1), subordinate (P2), and solitary (S) at the end of the experiment based on their emergent social positions. B) *Feeding behavior scoring method for pairs and solitary fish*. During feeding, pairs were separated by a divider to prevent competition, divider was included in solitary tanks as a sham control, and individuals were provided with equal amounts of pre-portioned food daily. Feeding behavior was quantified weekly from video recordings by counting the number of bites taken by each fish over a 5-minute period starting from the first bite. Initially, paired fish were denoted as P-max and P-min based on the total number of bites taken, because individuals could not be reliably identified due to their similar size and lack of distinguishing markings. From week 3 onwards, bites taken were scored separately for dominant (P1) and subordinate (P2) fish, in pairs where fish could be reliably identified based on clear size differences. C) *Agonistic behavior scoring method for pairs.* Agonistic behaviors (aggression, submission) were quantified during a five-minute observation period from video recordings of paired interactions. Aggression was quantified by the number of chases, headbutts, and bites, while submissive behavior included flees and shakes. Initially, total aggression and submission were measured at the tank level, because fish were similar in size and lacked distinguishing markings. From week 3 onward, agonistic behaviors were scored separately for dominant (P1) and subordinate (P2) fish, in pairs where individuals could be reliably identified due to the emergence of clear size differences. In the accompanying illustrations, the focal fish is shown in orange, with its partner depicted in black and white. D) *Gene expression profiling for pairs and solitary fish.* At the end of the experiment (week 5), whole-body samples were collected from a subset of individuals [n = 15 per social role: dominant (P1), subordinate (P2), solitary (S)]. To maximize detection of gene expression patterns associated with social roles, replicates with the largest size differences between dominant and subordinate fish were selected for transcriptomic profiling. The resulting RNA-seq data were used to compare molecular signatures across emergent social roles. In all panels, the timeline at the top indicates when (week 1 – week 5) and how (calipers, videos, tissue samples) data was collected.

To test these predictions, we compared the four phenotypic aspects of juveniles housed as size-matched pairs and size-matched solitary individuals. This experimental design allowed individuals equal opportunity to attempt taking on the dominant social role, however, through continuous social interactions (or in the case of solitaries, due to lack of social interactions), individuals took on varying social roles as dominant and subordinate members. At the end of the experiment, individual identities within pairs were verified using gene expression data or microsatellite genotyping (see Supplementary Materials), and the larger individual in each pair was denoted as pair rank 1 (P1), the smaller as pair rank 2 (P2), and solitary individuals as S (Fig. 1A). After excluding three replicates due to mortality, 27 replicates were used for downstream analysis (N = 54 paired fish, N = 27 solitary fish). Given that all phenotypes are entirely dependent on social context with no intrinsic baseline in this species, we arbitrarily chose P1 (dominant) as the statistical reference group for all downstream analyses. Standard length measurements taken at the beginning and end of the experiment were used to calculate growth over five weeks (Fig. 1A). Feeding behavior (Fig. 1B), and agonistic behaviors (aggression, submission; Fig. 1C) were measured weekly over five weeks, and whole-body gene expression was profiled after week five in a subset of 15 replicates (n = 45 fish; 15 P1, 15 P2, 15 S; Fig. 1D) selected to represent distinct growth trajectories across social roles. Our results show that an individual’s social context shapes its growth trajectory through coordinated changes in feeding behavior, agonistic behaviors, and gene expression: dominant individuals grow faster, eat more, exhibit more aggression, and upregulate genes linked to growth and ossification while suppressing satiety signals, compared to subordinate and solitary fish.

## Results

### Dominant individuals exhibit accelerated growth compared to subordinates and solitaries

To test the prediction that paired size-matched rivals will grow faster than size-matched solitary individuals, and one of the pair will grow faster than the other, as they attempt to outgrow each other, standard length (SL) was measured at the beginning and end of the five-week experiment (Fig. 1A). A generalized linear mixed model (GLMM) revealed a significant effect of time (DF = 1, F-value = 159.794, p < 0.0001), social position of fish (DF = 2, F-value = 31.030, p < 0.0001), and their interaction (DF = 2, F-value = 10.534, p < 0.0001) on fish size. At the beginning of the experiment (t1), post-hoc Tukey tests revealed that there was no difference in the size of P1, P2, or S (all P > 0.05; Fig. 2A), consistent with the initial size-matching design. At the end of the experiment (t5), P1 was significantly different from P2 and S (p < 0.001). Parameter estimates revealed that P1 individuals grew twice as much as P2 and S (Fig. 2A.1). The model, including the fixed effects (social position and time) and random effect (replicate ID), explained 65% of the variation in size (R^2^c = .650, R^2^m = .527; Fig. 2A).

**Figure 2:**
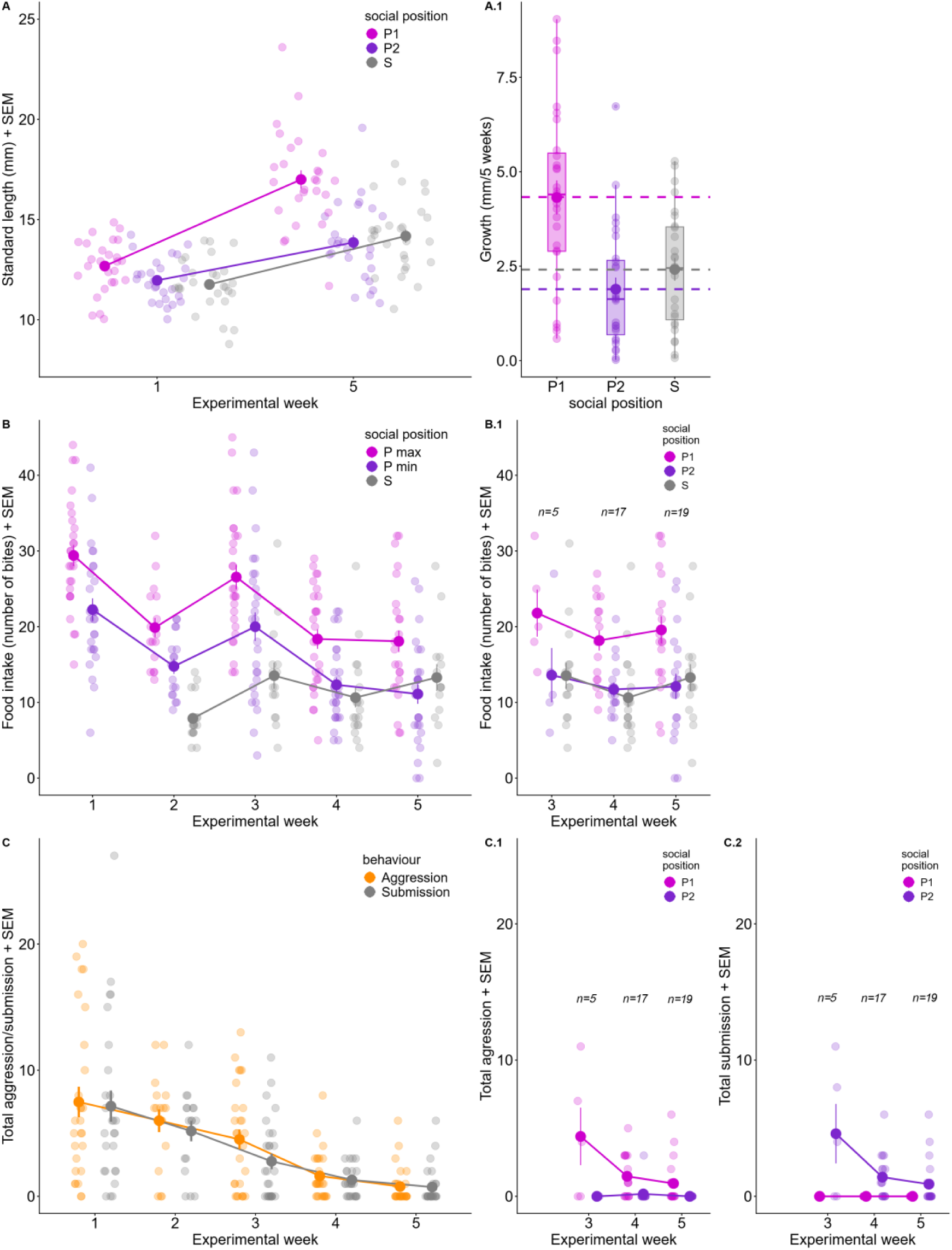
**A) Growth over time.** Standard length (mm) of individuals of each social position (pair rank 1 = P1; pair rank 2 = P2; solitary = S) over time with mean and standard error of mean (SEM). **A.1)** Overall growth (mm) over 5 weeks of each social position are shown with mean (dashed lines) and SEM. **B) Feeding behavior over time.** Individual food intake, measured in number of bites of size-matched pairs (P-max and P-min) vs solitary (S) individuals over time with mean and SEM. **B.1)** Food intake for a subset of replicates in which dominant (P1), subordinate (P2), and solitary (S) individuals could be reliably identified due to the emergence of clear size differences (week 3, n = 5; week 4, n = 17; week 5, n = 19), shown over time with mean ± SEM. **C) Agonistic behavior of pairs over time.** Aggression (the total number of chases, bites, and headbutts) and submission (based on the total number of flees and shakes) within pairs over time with mean and SEM. **C.1)** Aggression and **C.2)** Submission for a subset of pairs in which dominant (P1) and subordinate (P2) individuals could be reliably identified based on clear size differences (week 3, n = 5; week 4, n = 17; week 5, n = 19), shown over time with mean and SEM. Data points shown jittered on all figures for visual aid, but collected at the same time.

### Higher food intake in paired rivals during periods of intense conflict over size

To test the prediction that feeding behavior will be dependent on social position, and that paired size-matched rivals will eat more than solitary individuals as they attempt to outgrow each other, we quantified food intake. The number of bites of food taken by each fish following the first bite was scored from videos, across five weekly timepoints (Fig. 1B). Due to time constraints, no videos were recorded for solitary fish during week 1. All fish were provided equal quantities of food, and paired individuals were separated during feeding to prevent interference. A divider was also placed in solitary tanks during feeding as sham control. Given that P1 and P2 identities were not initially distinguishable in videos, individuals were classified based on feeding behavior within pairs as P-max (higher intake) or P-min (lower intake) while solitaries were denoted as S. The best-fit GLMM revealed a significant effect of time (experimental week; DF = 4, F-value = 27.6042, p < 0.0001), social position (P-max, P-min, S; DF = 2, F-value = 58.3621, p < 0.0001), and their interaction (DF = 7, F-value = 2.2745, p = 0.0288) on feeding behavior. Post-hoc pairwise contrasts revealed that both P-max and P-min ate more than S early in the establishment of the social hierarchy (weeks 1-3) (all p < 0.01); however, no significant difference was found between P-min and S during weeks 4 and 5 (both p > 0.05; Fig. 2B). Significantly higher food intake was found when comparing P-max and S during weeks 4 (p < 0.01), however, no significant difference was found by week 5 (p > 0.05). Within pairs, contrasts revealed significant differences between P-max and P-min throughout the experiment (all p < 0.05), except week 2 (p = 0.776). The model, including the fixed effects (social position and time) and random effect (replicate ID), explained 55% of the variation in feeding behavior (R^2^c = .552, R^2^m = .445; Fig. 2B).

For a subset of replicates in which dominant (P1) and subordinate (P2) members could be distinguished due to the emergence of clear size differences (week 3, n=5; week 4, n=17; week 5, n=19; Fig. 2B.1), we performed a complementary analysis using known social roles (P1, P2, S rather than P-max, P-min, S) to test the prediction that feeding behavior will be dependent on social position. In this subset, the best-fit GLMM revealed a significant effect of social position (P1, P2, S; DF = 2, F-value = 29.706, p < 0.0001), but not time (weeks 3–5; p=0.104) on feeding behavior. Post-hoc comparisons showed that P1 consistently ate more food than both P2 and solitaries (all p < 0.0001) in these later weeks, while no significant differences were observed between P2 and solitaries (p = 0.882). The model, including fixed effect of social position and random effect of replicate ID, explained 63% of the variation in feeding behavior (R^2^c = .627, R^2^m = .233; Fig. 2B.1).

### Agonistic behaviors (aggression, submission) decline as social roles stabilize

To test the prediction that in paired, size-matched rivals, agonistic behaviors will decline over time as size differences emerge, we scored aggression and submission from weekly videos (Fig. 1C). Aggression was quantified based on the sum of all bites, chases, and headbutts within pairs, while submission was quantified by the total numbers of flees and shakes (Wong et al., 2013, 2016). Because individuals within pairs could not be reliably distinguished on video, scores reflect total agonistic behavior per pair rather than per individual; however, aggression was typically only displayed by one individual while submission was typically only displayed by the other. Aggression and submission were not observed in solitary fish. The GLMM revealed that experimental week was a significant predictor of total aggression (DF = 4, F-value = 16.073, p < 0.001). Post-hoc pairwise comparisons showed significant differences in total aggression among most non-consecutive weeks (most p < 0.05), e.g., week 1 and 3, week 2 and 4, week 3 and 5 (Supplementary Table S3). Aggression within pairs declined over time (Fig. 2C). The model, including the fixed effect (time) and random effect (replicate ID), explained 42% of the variation in total aggression (R^2^c = .417, R^2^m = .306; Fig. 2C). Furthermore, the results showed experimental week was a significant predictor of total submission (DF = 4, F-value = 18.055, p < 0.001). As with aggression, Tukey’s post-hoc comparison showed significant differences in total submission among most non-consecutive week comparisons (most p *<* 0.05; Supplementary Table S3). The model, including the fixed effect (time) and random effect (replicate ID), explained 47% of the variation in total submission (R^2^c = .476, R^2^m = .308; Fig. 2C).

For a subset of pairs in which dominant (P1) and subordinate (P2) members could be reliably identified due to the emergence of clear size differences (week 3, n=5; week 4, n=17; week 5, n=19; Fig. 2C.1 & 2C.2), we examined how aggression and submission varied with known social roles (P1, P2). Using these distinctions, the best-fit GLMM showed that time (weeks 3–5; DF = 2, F-value = 3.883, p = 0.02), social position (P1, P2; DF = 1, F-value = 43.478, p < 0.0001), and their interaction (DF = 2, F-value = 7.455, p = 0.001) significantly predicted total aggression. Post-hoc contrasts revealed a decrease in aggression over time for P1 individuals, with significant differences between weeks 3–4 (p = 0.0009) and 3–5 (p = 0.0001), but not 4–5 because aggression in both weeks was close to zero (p = 0.477). No significant changes occurred for P2 fish whose aggression in all weeks was close to zero (p > 0.05). This model explained 52% of variation in total aggression (R²c = 0.522, R²m = 0.311; Fig. 2C.2). Similarly, the best-fit GLMM for total submission showed significant effects of time (weeks 3–5; DF = 2, F-value = 4.675, p = 0.012), social position (DF = 1, F-value = 41.071, p < 0.0001), and their interaction (DF = 2, F-value = 7.374, p < 0.001). Post-hoc contrasts indicated decreased submission over time for P2 individuals, with significant differences between weeks 3–4 (p = 0.0004) and 3–5 (p < 0.0001), but not 4–5 because submission in both weeks was close to zero (p = 0.526). No significant differences were found for P1 individuals whose submission was always close to zero (p > 0.1). This model explained 48% of variation in total submission (R²c = 0.486, R²m = 0.332; Fig. 2C.3).

### Social roles shape global gene expression patterns

To test the prediction that whole-body gene expression patterns will vary with social roles once clear social positions have emerged (P1, P2 and S), we performed gene expression profiling on 45 individual fish (experimental groups containing paired individuals and corresponding solitaries; N = 15 samples per social position; Fig. 1D). TagSeq libraries (Meyer et al., 2011) were sequenced on NovaSeq 6000 (single end, 100 bp), yielding an average 8.2 million raw reads per sample (Supplementary Table S1). After quality filtering, reads were mapped to the *A. percula* genome (Lehmann et al., 2019), using STAR (Dobin et al., 2013), and gene-level counts were generated with *featureCounts*. Genes with a mean count <10 were excluded from downstream analysis. Differential gene expression analysis was performed using DESeq2 (Love et al., 2014), with the model ∼ social position + genotype (clutch ID), and genes were considered differentially expressed at an FDR adjusted p < 0.05. To assess global gene expression differences across social positions, count data were rlog-transformed and used for principal component analysis (PCA) based on Euclidean distances using the vegan package (Oksanen et al., 2017). PCAs were performed on all genes, and the top 50%, top 10%, and top 5% of the most variable genes, all of which showed consistent clustering across all pairwise PC comparisons (PC1 vs PC2; PC2 vs PC3; PC1 vs PC3; Supplementary Fig. S3). Of all the examined PCAs, PC2 and PC3 with all genes showed the clearest clustering by social position (p = 0.003), and revealed overall no significant difference in gene expression between P2 and S individuals, with P1 individuals exhibiting more distinct clustering (Fig. 3A; replicate-annotated version of Fig. 3A in Supplementary Fig. S4; for all other PCAs see Supplementary Fig. S3). Pairwise contrasts across social position further confirmed no differentially expressed genes (DEGs) between P2 and S individuals (Fig. 3B; Supplementary Fig. S5). A total of 96 DEGs were significantly upregulated in P2 individuals, and 390 DEGs were significantly upregulated in S individuals, when compared to P1. Eighty-four of these upregulated DEGs were shared between P2 and S (Supplementary Table S4). A total of 37 DEGs were significantly downregulated in P2 individuals, and 130 DEGs in S individuals, when compared to P1 individuals. A total of 25 downregulated DEGs were shared between P2 and S individuals (Fig. 3B; Supplementary Table S4).

**Figure 3:**
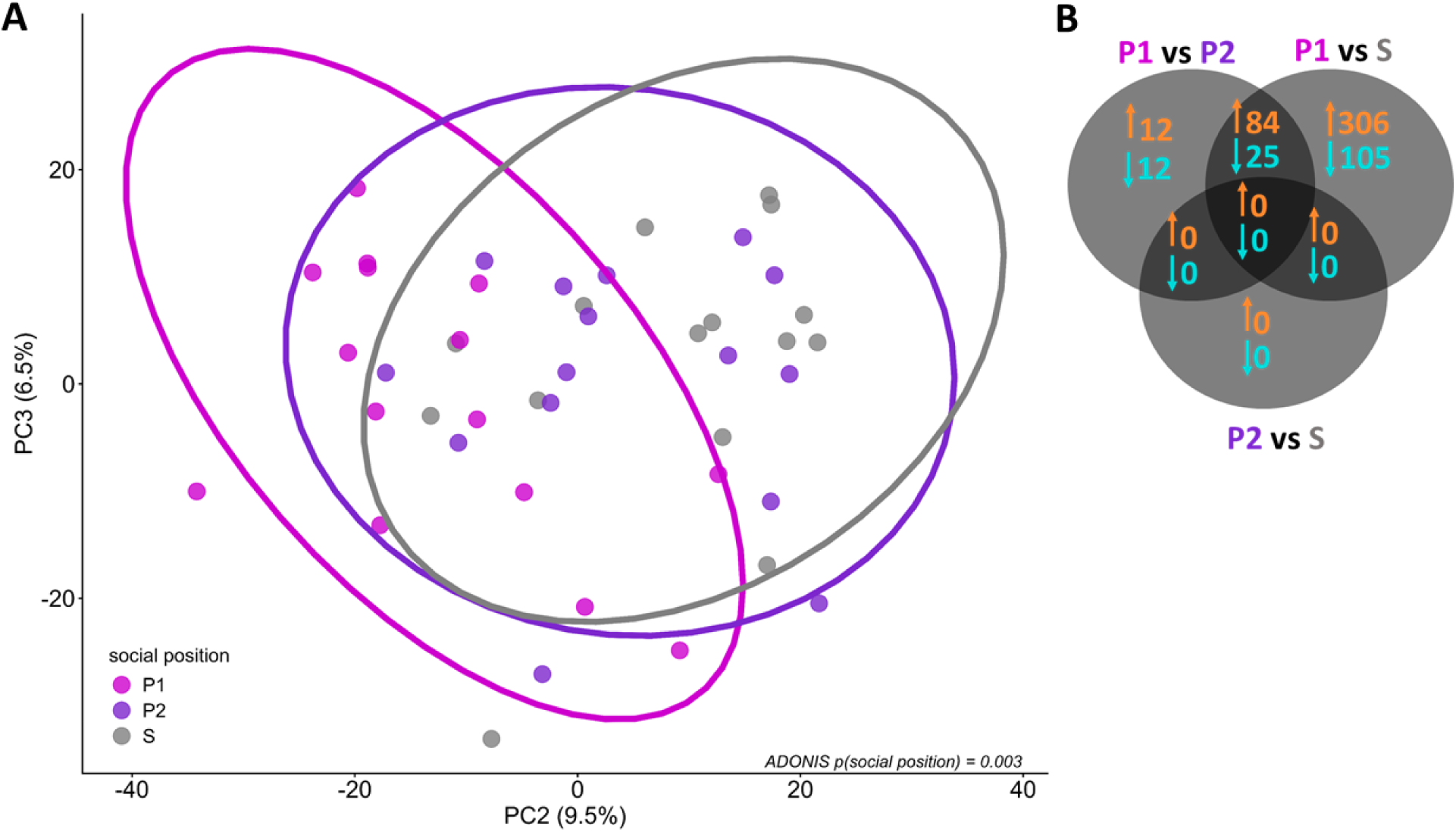
Overall gene expression profiles showed no significant differences between P2 and S, compared to P1 individuals. **A)** Principal Component Analysis (PCA) of all genes across social positions, accounting for clutch ID (genetic background), with PERMANOVA results showing a significant effect of social position. Ellipses show the estimated confidence intervals from a 95% multivariate t-distribution. **B)** Venn diagram illustrating differentially expressed genes (DEGs) from pairwise comparisons of social positions (FDR-adjusted p-value of 0.05). Orange upward arrows and corresponding numbers represent significantly upregulated DEGs in the first term, relative to the second, while blue downward arrows and numbers indicate downregulated DEGs (Supplementary Table S4).

### Gene co-expression networks associate expression patterns with phenotypic traits

To test the prediction that global gene expression patterns will be correlated with observed phenotypic changes in growth and feeding behavior while accounting for the individual’s social position, a weighted gene co-expression network analysis (WGCNA; Langfelder & Horvath, 2008) was conducted on the rlog-transformed gene expression (GE) dataset. Unlike pairwise DEG analysis, which treats genes independently, WGCNA identifies groups of co-expressed genes (modules) and tests whether modules of eigengene expression are correlated with variation in phenotypes of interest. For modules showing significant correlations with phenotypes, gene ontology (GO) analyses were performed using Fisher’s exact tests to identify over-represented pathways within each module. A total of 9 WGCNA modules were found (Supplementary Fig. S6; only 8 modules are shown, as the 9th contained over 10,000 genes with no clear eigengene expression pattern and was therefore excluded). Of the remaining modules, four (brown, blue, red, black) showed significant associations with the phenotypes of interest and were explored by GO analyses (Supplementary Fig. S7).

### Growth related genes are upregulated in dominant individuals

The brown module identified via WGCNA showed strong positive correlations with body size (Supplementary Fig. S6). Subsequent GO enrichment analysis revealed enrichment for biological process (BP) and molecular function (MF) GO terms associated with development, ossification, morphogenesis, and cytoskeletal regulation (Fig. 4). From the genes associated with these GO terms, a total of 38 genes were significantly upregulated in P1 individuals compared to P2 and S fish (FDR p ≤ 0.05; Fig. 5A). These included extracellular matrix and bone development genes (*col1a2, col6a1, mmp14b, plod2, loxa*), signaling factors (*bmp4, tgfb1a, tgfb3, fstl1b*), and regulators of muscle and cytoskeletal function (*myh7ba, tpm1, actn2b, mstna, smo*, among others; Fig. 5B).

**Figure 4:**
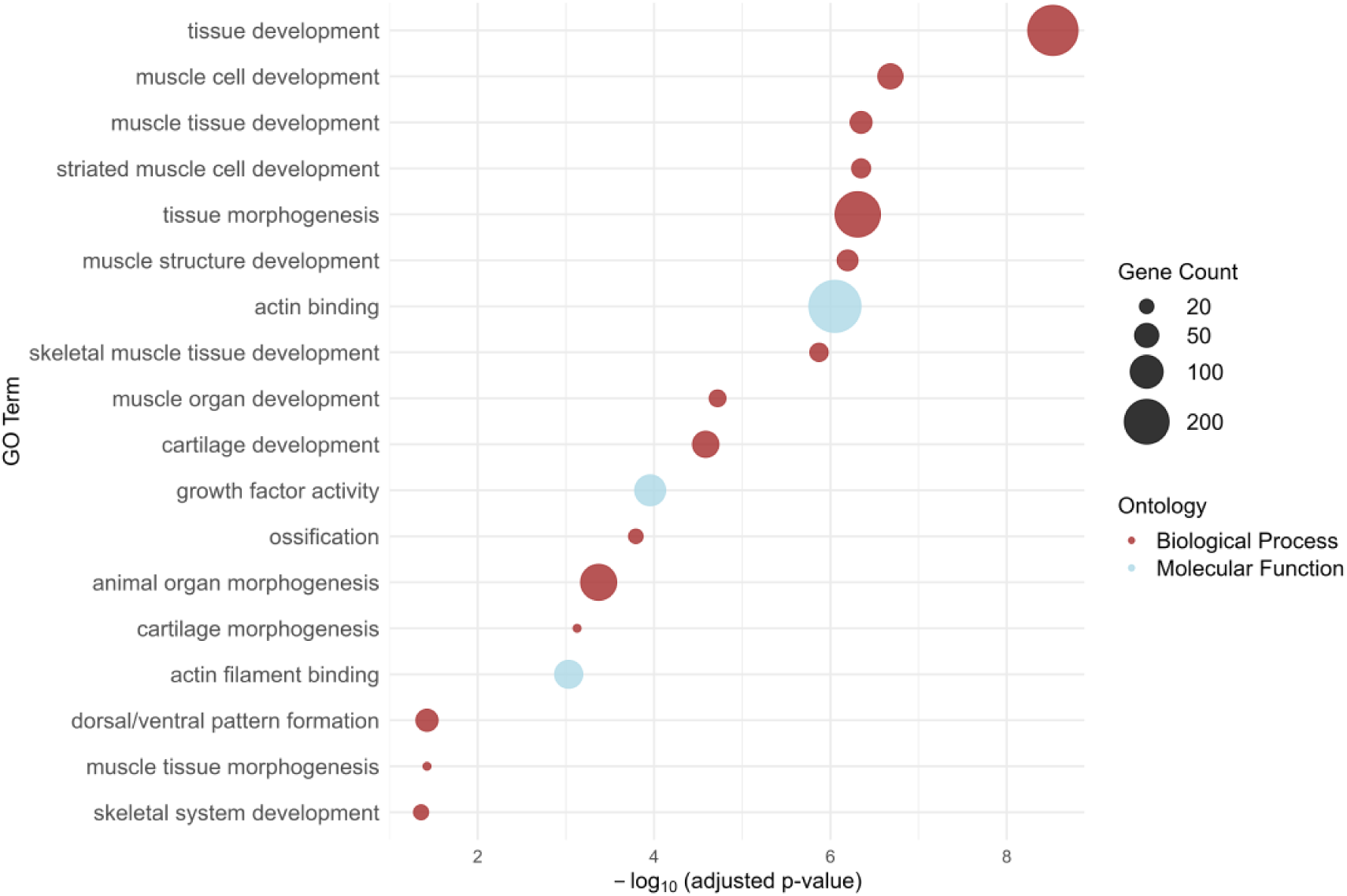
Gene ontology (GO) enrichment of the brown module associated with growth and ossification. An exhaustive list of GO enrichment results for genes assigned to the brown WGCNA module (Fig. 5). Bubble size reflects numbers of genes associated with each term, color indicates GO category. Terms are ordered by adjusted p-value significance on the x-axis (–log₁₀ scale), with more significant terms on the right.

**Figure 5:**
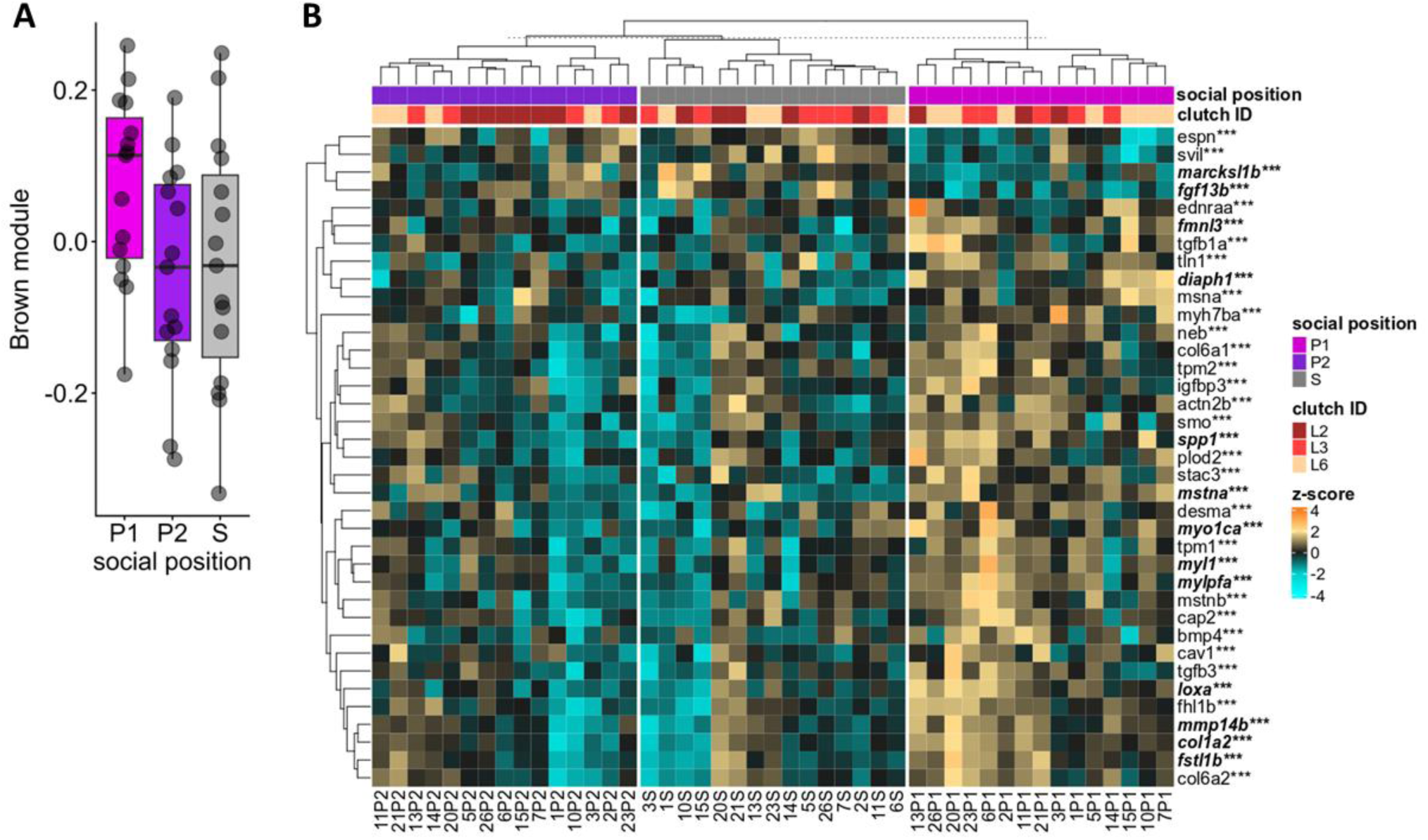
P1 individuals showed upregulation of genes associated with growth and ossification. **A)** Average eigengene values across social positions for the brown module. **B)** Heatmap of significant DEGs (FDR p-value ≤ 0.05) of growth and ossification genes in the brown Biological Process (BP) and Molecular Function (MF) GO terms across the three social positions. Samples are grouped *a priori* by social position (dominant, P1; subordinate, P2; solitary, S) and ordered within each group by column dendrograms reflecting expression similarity within social positions. The row dendrograms show relatedness among genes based on expression profile similarity across all samples. Genes that are significantly differentially expressed between individuals of P1 vs P2 are denoted with an asterisk (*), those between P1 vs S with a double asterisk (**), and three asterisks (***) represent both comparisons. Genes shown in ***bold and italics*** indicate significance at an FDR adjusted p-value < 0.1.

### Dominant social role is associated with suppression of satiety signals and altered metabolic pathway activity

While the red WGCNA module was positively associated with appetite-related gene expression, we found no strong enrichment of appetite and metabolism related GO terms (Supplementary Fig. S7). To further examine differences in appetite and metabolism across social roles, we analyzed genes known to play a role in these processes identified in the sister species *Amphiprion ocellaris* (Herrera et al., 2025). High-confidence *A. percula* orthologs were identified using a reciprocal BLAST+ approach (Camacho et al., 2009) (Supplementary Table S2) and matched to our whole-body expression dataset. We found a set of genes controlling appetite (*ghrl, insra, irs2a, irs2b, adycap1b, cckb*) to be significantly downregulated in P1 individuals compared to P2 and S (FDR p ≤ 0.01; Fig. 6). Similarly, genes that control energy metabolism via TCA cycle (*mdh1aa, suclg1, suclg2, sdha, ogdhb, ogdhl*), and glycolysis (*gapdhb, aldob, tpi1a, eno2b*) were also significantly downregulated in P1 individuals compared P2 and S individuals (Fig. 6).

**Figure 6:**
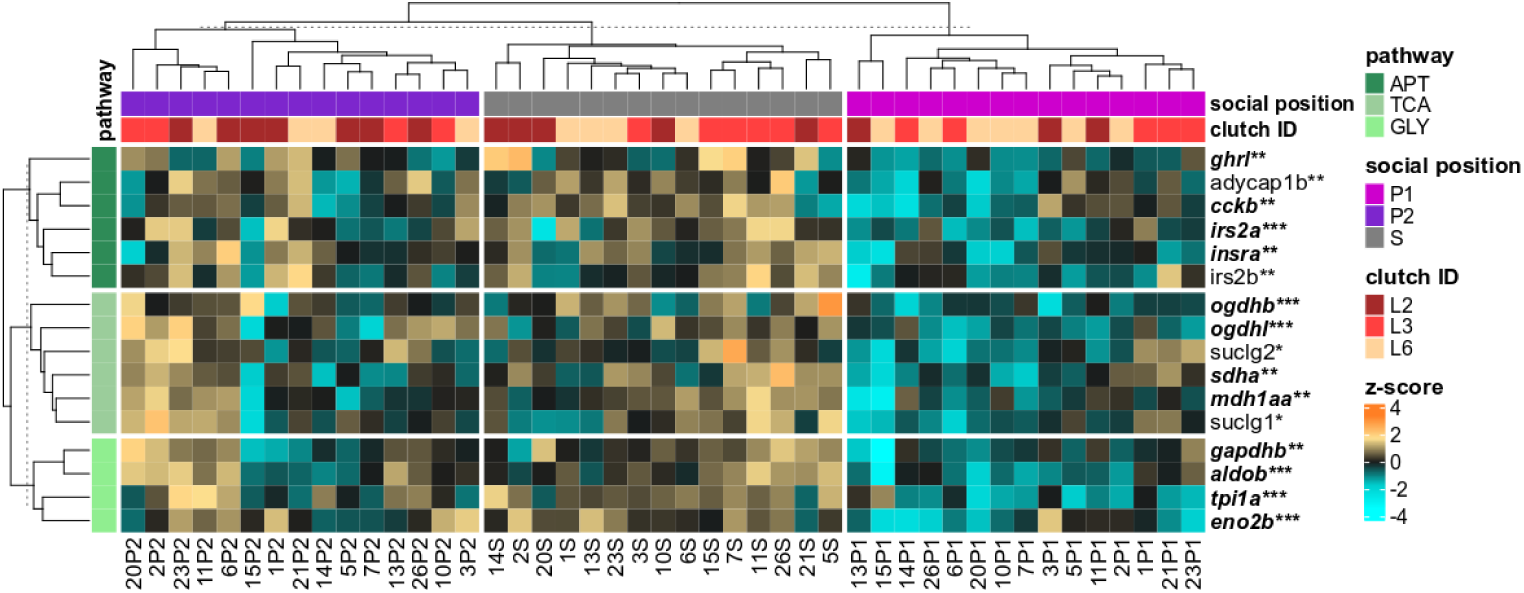
P1 individuals showed downregulation of metabolic and appetite suppression genes. Heatmap of significantly differentially expressed genes (DEGs) (FDR p-value ≤ 0.01) of genes associated with appetite regulation (APT), Krebs cycle (TCA), and glycolysis (GLY) pathways across the three social positions indicating genetic background of individuals (clutch ID). Samples (columns) are grouped *a priori* by social position (dominant, P1; subordinate, P2; solitary, S) and ordered within each group by the column dendrogram, which reflects expression similarity among samples within each social position. Genes (rows) are grouped *a priori* by pathway (APT, TCA, or GLY), and within each pathway group, the row dendrogram reflects relatedness among genes based on expression profile similarity across all samples. Genes that are significantly differentially expressed between individuals of P1 vs P2 are denoted with an asterisk (*), those between P1 vs S with a double asterisk (**), and three asterisks (***) represent both comparisons. Genes shown in ***bold and italics*** indicate significance at an FDR adjusted p-value < 0.1.

## Discussion

Here we show that in paired, size-matched rivals, individual growth trajectories are associated with coordinated changes in feeding behavior, agonistic behaviors, and underlying gene expression, leading to the emergence of clear social roles. We found that the individual ultimately emerging as P1 (dominant) grew significantly more compared to individuals taking on the P2 (subordinate) role. Moreover, S (solitary) individuals showed similar growth to P2 fish (Fig. 2A). To facilitate this differential growth dependent on social position, we have shown that early in the experiment both paired individuals, P-max and P-min, ate more per capita compared to S individuals (Fig. 2B). Later in the experiment, as size differences within pairs emerged, P1 ate more than either P2 or S (Fig. 2B.1). In conjunction with increased food intake in pairs, we have also shown that agonistic behaviors within pairs were significantly higher at the beginning of the experiment compared to the end of the experiment, once size differences were established (Fig. 2C). Furthermore, aggressive behaviors were displayed by the dominant members of pairs while submissive behaviors were shown by subordinate fish (Fig. 2C.1 & 2C.2). These findings show that coordinated changes in growth, feeding behavior, and agonistic behaviors reinforce phenotypes that match each individual’s social role. Interestingly, in 25% of the 27 replicates, the initially smaller fish ultimately emerged as the dominant member, demonstrating that winners and losers were determined during early social interactions rather than being fully predetermined by initial size differences. These behavioral and morphological shifts were mirrored at the molecular level. P1 individuals showed significantly different gene expression profiles compared to P2 and S individuals with P1 individuals upregulating genes linked to ossification (e.g., *collagen, myoglobin, bmp, and sost*) relative to P2 and S fish (Fig. 5). Likewise, genes that promote satiety signals (e.g., *cholecystokinin (CCK), adycap1b*) were downregulated in P1 relative to P2 and S, consistent with their heightened appetite, suggesting appetite suppression in P2 and S individuals (Fig. 6). Additionally, metabolic pathway genes involved in the TCA cycle (*ogdhl, ogdhb, sdha, mdh1aa*) and glycolysis (*tpi1a, eno2b, gapdhb, aldob*) were more active in P2 and S fish compared to P1 (Fig. 6), suggesting different energy utilization strategies.

Our findings on growth across social positions are broadly consistent with previous research, showing that size-matched rivals exhibit divergent growth trajectories based on their ultimate social rank. P1 individuals grew significantly more than P2 (mean ± SD: 4.32 ± 2.34 mm vs. 1.89 ± 1.60 mm), consistent with earlier studies in clownfish (Desrochers et al., 2020; Reed et al., 2019) and other social vertebrates such as meerkats (Huchard et al., 2016). Desrochers et al. (2020) also observed that P1 individuals grew 5.0 mm on average, compared to 3.1 mm for P2s and 1.6 mm for solitary fish, in a similar time frame. Consistent with previous findings, our P1 individuals exhibited significantly greater growth (4.32 ± 2.34 mm) than P2 individuals (1.89 ± 1.60 mm) or S individuals (2.41 ± 1.55 mm). In contrast to previous findings, we did not detect a difference between P2 (1.89 ± 1.60 mm) and S individuals (2.41 ± 1.55 mm). This contrast is likely due to improved experimental conditions in our study, particularly by the inclusion of host anemones, which seemed to improve the growth and condition of solitary individuals relative to previous experiments (Desrochers et al., 2020; Reed et al., 2019; Rueger et al., 2022).

Furthermore, this is amongst the first studies where the adjustment of feeding behavior across social positions has been quantified in clownfish. Despite equal food availability provided to all individuals, we show that feeding behavior differed significantly across social ranks, with individuals adjusting their food intake in accordance with their social roles. Similar socially mediated regulation of food intake has been reported in Kalahari meerkats (Huchard et al., 2016) and emerald coral goby (Wong et al., 2008). Meerkat size-matched rivals closely monitor each other’s food intake and modulate their own in an attempt to match growth and body size (Huchard et al., 2016), while emerald coral goby subordinates have been shown to “diet” as they near their immediate dominant’s size, to avoid conflict (Wong et al., 2008). Our findings suggest a comparable mechanism in clownfish, wherein feeding behavior is fine-tuned to emerging social roles. Additionally, here we quantified aggression and submission within pairs during the establishment of the dominance hierarchy, revealing peak agonistic interactions during the first two weeks of the experiment. This is consistent with other evidence showing that conflict intensity increases as size differences between individuals decrease, with the highest level of conflict observed between size-matched rivals (Bender et al., 2005; Wong et al., 2016). Within pairs, we show that conflict over size is resolved by taking on differing social roles via the adjustment of growth trajectories, feeding behavior, and agonistic behaviors. Indeed week 3 onward, when pair members could be reliably identified, dominant individuals consistently consumed more food and displayed aggressive behaviors, while subordinates ate less and exhibited submissive behaviors, clearly reflecting the emergence of two distinct social roles. While we cannot be certain, it is highly likely that these differences were present during the earlier weeks as well. Likely, winners and losers emerge soon after their initial interactions (Milewski et al., 2022; Wong et al., 2016), allowing members of the social group to adjust their social role while progressing into their social position.

In vertebrates, growth is regulated by four major signaling pathways: the Growth Hormone (GH)/IGF pathway, the insulin/PI3K-AKT pathway, the mTOR pathway, and the Hippo pathway (Fuentes et al., 2013; Glass, 2003). Together, these pathways coordinate cellular processes that promote overall growth and ossification. Their downstream effects are reflected in several functional gene categories, including those involved in ossification, bone matrix formation and structure, muscle development and contraction, and other structural regulators (Valenti et al., 2020). P1 individuals exhibited significantly greater growth than P2 and S fish, which was accompanied by coordinated upregulation of genes involved in skeletal and muscular development. This growth phenotype appears to be regulated via conserved vertebrate hormonal pathways. We found genes associated with the GH/IGF axis, including *igsbp3*, a proliferation-promoting member of the IGF-binding protein family (Zheng et al., 2024), and *mstna* and *mstnb*, negative regulators of muscle growth (Lee & McPherron, 2001), upregulated in P1 individuals, consistent with a growth-promoting state. For the insulin/ PI3K-AKT pathway we showed significant upregulation of insulin receptor genes (e.g., *insra, irs2a, irs2b*) and several structural and cytoskeletal genes known to be downstream targets of PI3K-AKT signaling (*cap2, tln1, actn2b*) (Glass, 2010). The mTOR pathway, which drives protein synthesis and cellular hypertrophy (Schiaffino et al., 2013), was also found to be activated as P1 individuals showed upregulation for muscle contractile genes (*myh7ba, myl1, mylpfa, tpm1, tpm2*) and cytoskeletal regulators (*diaph1, fmnl3*) (Gunning et al., 2008). Additionally, increased expression of *smo*, a Hedgehog pathway gene that interacts with Hippo signaling, points to active control of tissue growth boundaries (Kodaka & Hata, 2015). These four pathways appear to stimulate structural and matrix-related genes: P1 fish upregulated bone matrix collagens (*col1a2, col6a1, col6a2*), matrix remodeling enzymes (*mmp14b, plod2, loxa*), and morphogenetic regulators (*bmp4, tgfb1a, tgfb3, fstl1b*), all contributing to bone growth and extracellular matrix turnover (Gistelinck et al., 2016; van Boxtel et al., 2011). A key mineralization gene, *ssp1*, was also upregulated in P1 fish, supporting the activation of osteogenic pathways (Windhausen et al., 2015). In contrast, P2 and S individuals showed consistent downregulation of these genes and pathways, aligning with their reduced growth. These results provide a broad framework for understanding the molecular mechanisms of strategic growth in other social vertebrates. Pathways underlying skeletal and muscular morphogenesis are highly conserved between mammals and fish (Howe et al., 2013; Valenti et al., 2020), suggesting that similar molecular mechanisms may underlie skeletal remodeling during rank transitions of meerkats and mole-rats. Nonetheless, some differences in these pathways are expected, particularly the capacity for bone regeneration, which is present in fish but absent in mammals (Pfefferli & Jaźwińska, 2015).

In our study, P1 individuals showed increased food intake compared to P2 individuals. To explore underlying GE differences, we examined genes known to be associated with appetite regulation and metabolism in *A. ocellaris* (Herrera et al., 2025), and found downregulation of these genes in P1 individuals compared to P2 and S. As time since last meal can strongly alter GE profiles, all fish were fed ∼14 hours before euthanasia; however, residual pellets may have remained, making the exact time of last ingestion uncertain. To assess whether differences in appetite-related gene expression could underlie the distinct feeding behaviors associated with dominant and subordinate roles, we explored appetite suppressor genes (e.g.: cholecystokinin *(CCK),* cocaine- and amphetamine-regulated transcript *(CART),* leptin *(leptin),* peptide YY *(PYY), and* oxytocin *(OXT)*) that usually rise after feeding, as well as appetite stimulators (e.g.: ghrelin *(ghrl),* agouti-related neuropeptide *(AGRP),* neuropeptide Y *(NPY),* and hypocretin neuropeptide precursor *(HCRT*)), that typically increase during fasting to promote feeding behavior (Volkoff, 2019). We found two appetite suppressor genes, *CCK* and *adycap1b*, to be significantly downregulated in P1 individuals. *CCK* functions as a short-term satiety signal, increasing after feeding and decreasing during fasting in goldfish (Thavanathan & Volkoff, 2006). Similarly, *adycap1b*, a homolog of *PACAP2*, has been shown to suppress food intake in zebrafish (Nakamachi et al., 2019). Of the appetite stimulator genes, we observed downregulation of *ghrl* in P1 individuals compared to P2 and S. Ghrelin is traditionally classified as an appetite stimulator, typically increasing during fasting to promote feeding (Assan et al., 2021; Soengas, 2021). However, in teleosts, ghrelin can also act as a short-term satiety signal (Volkoff, 2019). The observed concurrent downregulation of *CCK*, *adycap1b* and *ghrl* of P1 individuals points towards a reduced need to suppress feeding, providing a potential mechanistic explanation for higher food intake in dominant fish, which reinforces their social role. A similar pattern has been observed in domesticated chickens, where rapid growth is associated with increased appetite, which is facilitated via the decreased expression of *CCK*, leading to an altered satiety signal (Dunn et al., 2013). Molecular signals indicative of appetite suppression were likely even more pronounced earlier in the experiment (e.g., at week 2), when we showed increased food intake coinciding with intense conflict over size.

Another way of thinking about these results, since all of the gene expression measures are relative and there is no intrinsic baseline in this species, is to take the perspective of P2 individuals: P2 individuals exhibited lower food intake and upregulation of appetite-related genes associated with satiety. Upregulation of appetite suppressor genes (*CCK and adycap1b*) may reflect a physiological drive to suppress appetite. This pattern is consistent with the observed subordinate strategy of reduced food intake, providing a molecular basis for social position driven behavioral restraint. Indeed, at the molecular level this reduced food intake might parallel the “dieting” behavior observed in emerald coral goby subordinates, a strategy thought to limit growth and reduce conflict with immediate dominants (Wong et al., 2008). However, this behavioral strategy in the context of dominance hierarchies has not been demonstrated at the molecular level. *Ghrl* was also significantly upregulated in P2 individuals, despite their lower food intake. In rainbow trout, it has been shown that ghrelin-treated individuals exhibited reduced appetite, ultimately leading to a ghrelin-induced decrease in growth (Jönsson et al., 2010). Interestingly, increased ghrelin expression was also shown to be associated with increased locomotion both in goldfish and zebrafish (Matsuda et al., 2006; Navarro-Guillén et al., 2019). Given that P2 fish tend to be subjected to aggressive behaviors in the form of chases, increased *ghrl* levels may support the need for increased movement.

From a metabolic perspective, P1 individuals also exhibited downregulation of genes involved with aerobic metabolism (TCA cycle; *ogdhl, ogdhb, sdha, mdh1aa* genes) and anaerobic glycolysis (*tpi1a, eno2b, gapdhb, aldob* genes). This pattern is somewhat contradictory to previous findings (Lu et al., 2020; Roux et al., 2023; Yin et al., 2023), and unexpected given the increased growth observed in P1 individuals, a process that is energetically costly and dependent on ATP production. Some parallels can be drawn with freshwater cavefish, where fast-growing individuals also showed downregulation of some key glycolysis genes (*aldob, eno2*) compared to slow-growing counterparts (Yin et al., 2023), however, they found other glycolysis genes (e.g.:*agl, pfkm, gapdh, bpgm*) to be upregulated in fast-growing fish suggesting a more nuanced regulation. It is possible that P1 individuals rely on glycolysis under conditions of high food availability to facilitate growth, but we failed to detect such signals as they are likely tissue specific and were not detectable via our whole-body RNA-seq approach. P1 individuals also showed downregulated insulin receptor genes (*insra, irs2a, irs2b*), which regulate glucose uptake and energy balance post-feeding (Sheridan, 2021). This pattern may be consistent with a shift toward fatty-acid β-oxidation metabolism, which yields substantially more ATP than glycolysis (Darvey, 1998) and could support increased growth. However, we did not find any upregulated β-oxidation related DEGs in P1 fish. The appetite and metabolic profiles of P1 individuals characterized by the downregulation of appetite suppressors along with suppressed expression of both aerobic TCA cycle and anaerobic glycolytic genes, is somewhat contradictory. It is likely that P1 individuals are meeting their energy demands efficiently via increased food intake and allocating energy towards growth, although the precise pathways remain unresolved in our data.

In contrast, P2 fish showed significant upregulation of both aerobic TCA cycle genes and anaerobic glycolysis genes, as well as upregulation of insulin receptor genes. The pronounced upregulation of appetite-suppressing genes, alongside the elevated ghrelin expression, suggests that P2 individuals may be experiencing an energy deficit, likely due to reduced food intake and increased locomotory activity. Correspondingly, while it is somewhat contradictory, the upregulation of both aerobic (TCA cycle) and anaerobic (glycolysis) metabolic genes suggests a physiological response to meet energy demands during “dieting” behaviors. Finally, S individuals maintained a stable appetite but showed an analogous gene expression profile of appetite and metabolic genes to P2 individuals. This may be linked to the stress of social isolation, which in larval zebrafish has been shown to alter feeding behavior and suppress food intake (Wee et al., 2022). Although many well-known appetite-regulating genes and metabolic pathway genes were not differentially expressed, the overall expression patterns suggest that individuals modulate their molecular phenotypes to reinforce their social roles as dominant, subordinate, and solitary members.

A striking finding of our study was the similarity in growth trajectories, feeding behavior, and overall gene expression profiles between subordinate (P2) and solitary (S) individuals. Although solitary individuals had the opportunity to assume the dominant (P1) role in their tanks, they instead adopted subordinate-like characteristics, indicating that social context, not opportunity alone, drives social role expression. Similar patterns have been anecdotally observed in the wild, where juvenile clownfish remained small for extended periods (over four months) following the loss of a dominant partner, only initiating changes in social role towards dominant characteristics upon the arrival of a new group member (personal observations of 8+ instances during two long-term independent studies by Pete Buston and Lili Vizer). It’s almost as if these fish are remaining dormant, until their future social position becomes clear. Comparable results have been reported in Damaraland mole rats, where subordinate helpers and solitaries were indistinguishable both on a morphological and on a gene expression level (Johnston et al., 2021; Thorley et al. 2018) and therefore were treated as one group of non-breeders for downstream analysis. This approach, however, overlooks the differing social contexts these individuals experience. Other studies in cichlids (Hesse & Thünken, 2014), zebrafish (Wee et al., 2022), and mice (Benfato et al., 2022) have shown that social isolation often results in decreased growth, increased corticoid levels, altered appetite, and behavioral changes. However, none of these studies considered changes associated with social isolation in the ecological context of social behavior and dominance hierarchies. Our results suggest that the presence of a social partner is essential for the establishment of social roles, likely because progression through the social hierarchy involves irreversible transitions in social roles (Buston, 2003, 2004b), a pattern also true for Damaraland mole rats (Johnston et al., 2021). Consequently, social role differentiation appears to require mutual recognition and reinforcement within a group context.

We have used clownfish (*Amphiprion percula*) as a model system to investigate the role of phenotypic plasticity during the establishment of dominance hierarchy. We showed that an individual’s social context shapes its growth trajectory through coordinated changes in behaviors and underlying gene expression, with dominant members strategically upregulating their growth, while subordinates and solitaries strategically downregulating their growth. This study lays the foundation for a broader framework to explore the mechanisms underpinning strategic growth in social vertebrates. Future research should focus on characterizing periods of strategic upregulation and downregulation of growth dependent on social context, with particular emphasis on gene expression signatures. In our current study, individuals within pairs were ultimately progressing towards rank 1 dominant female and rank 2 subordinate male roles. In other species known to exhibit strategic growth, such as cichlids (Maruska & Fernald, 2014; Renn et al., 2008), meerkats (Huchard et al., 2016; Russell et al., 2004), and mole rats (Dengler-Crish & Catania, 2007; Johnston et al., 2021; Thorley et al., 2018) strategic upregulation of growth cannot be disentangled from sexual maturation, due to the nature of their dominance hierarchy. However, clownfish offers an ideal system to disentangle growth-related transcriptional profiles from those associated with sexual maturation, as subordinate individuals of rank 3-6 exhibit strategic growth, without signs of sexual maturation (Buston, 2003; Buston & Clutton-Brock, 2022; Reed et al., 2019).

In this study, we used whole-body transcriptomics, which revealed overarching expression patterns, however, we acknowledge that this approach limited our ability to detect finer-scale signals. Obviously, this approach cannot resolve tissue-specific gene expression changes (Lu et al., 2020; Roux et al., 2023; Yin et al., 2023) which are critical for a complete understanding social role differentiation, such as adjustments of behavior, appetite, and growth (a list we consider non-exhaustive). To disentangle gene expression signatures associated with socially induced phenotypes such as the strategic up- and downregulation of growth, future studies should include sampling points aligned with the onset of these physiological shifts. Moreover, to understand underlying pathways involved, tissue specific transcriptomics will be essential. Sampling multiple tissues across multiple time points would disentangle nuanced regulatory processes underlying coordinate functional shifts leading to different social phenotypes of strategic growth. These include appetite regulation (orexigenic and anorexigenic signaling), metabolic pathways (e.g., glycolysis, lactic fermentation, TCA cycle, fatty acid β-oxidation), growth-regulating pathways (GH/IGF, insulin/PI3K-AKT, mTOR, and Hippo), and potential molecular signatures to varying levels of social stress. Additionally, while laboratory studies can provide controlled environments, complementary field-based studies will be crucial to validate laboratory results. Once candidate hormonal or molecular pathways are identified, their functional roles could be experimentally confirmed through targeted hormonal or CRISPR manipulation (Mitchell et al., 2021). Together, these approaches provide a powerful framework for uncovering the dynamic, context-dependent mechanisms that regulate strategic growth and coordinated changes associated with the establishment and maintenance of dominance hierarchies in social vertebrates.

## Methods

### Adult Broodstock

We conducted these experiments at Boston University (Boston, MA, USA) during the summer of 2022. All fish used in these experiments were reared from broodstock that were wild-caught as non-breeders (less than 30 mm in standard length) in Papua New Guinea and supplied by Quality Marine and Sea Dwelling Creatures. The removal of non-breeders is considered a sustainable practice because they do not directly influence the survival, growth, or reproduction of the breeders (Buston, 2004a). Laboratory adult broodstock were housed as sixty pairs each in their own 120 L tanks with sand bottom and *Entacmaea quadricolor* anemone habitat on rocks over a 15×15 cm ceramic tile. A detailed description of broodstock housing conditions can be found elsewhere (Pacaro et al., 2023; Schlatter et al., 2022).

### Experimental Juveniles

To obtain experimental juveniles, egg clutches of adult broodstock were monitored daily until day 8, when they were moved to closed saltwater larval rearing tanks (20 L). To stimulate hatching, egg clutches were gently aerated using an air stone placed directly below the clutch. Multiple egg clutches hatched simultaneously, and the larvae were reared to 30 days of age in their respective tanks. During rearing, a daily water change of 10 L was conducted, and abiotic conditions of pH 8.1 ± 0.1, temperature 26 ± 1° C and salinity 35 ± 2 ppt were maintained. Hatched larvae were fed rotifers (*Brachionus rotundiformis*) (15 rotifers ml^-1^) in water tinted with *Nannochloropsis* algae paste (Rotigreen Nanno, Reed Mariculture, USA) between days 0-8. From day 4 until 14, decapsulated *Artemia salina* nauplii (3 Artemia ml^-1^) were fed. From days 4 to 8, both rotifers and Artemia were fed simultaneously to allow the larvae to gradually adjust to the transition between food sources. Once the fish reached 14 days of age, TDO-B1 Chroma BOOST Granule (Reef Nutrition) fish pellets were fed in addition to Artemia. Food pellets are composed of 49% crude protein, 13% crude fat, 9.7% carbohydrates. Juvenile fish were reared to 30 days of age, after which individuals were photographed, measured, and moved to experimental housings. All fish used in this experiment were sexually immature juveniles (one-month-old at the beginning, two-month-old at the end), as clown anemonefish reach sexual maturity between the ages of one and two years old. Also, under ideal conditions, individuals can remain sexually immature for a decade or more (Buston 2004b; Buston & García, 2007). Therefore, in this study, the emergence of social roles and associated phenotypes do not reflect changes associated with sexual maturation.

### Experimental Housing

Experimental fish were housed in 3.5 L breeder boxes with a sea anemone (*Entacmaea quadricolor)*, with a mean anemone surface area of 16.263 cm^2^ ± 1.739. Breeder boxes were submerged in 120 L tanks, which were part of a re-circulating saltwater aquarium system. Flow through each 120 L tank was approximately 9 L h ^-1^ ± 0.5 L. Abiotic conditions were monitored regularly and maintained as constant as possible. pH (8.35 ± 0.45), temperature (26.7 ± 1.2°C) and salinity (34.5 ± 1.5 ppt) were monitored daily by GHL profilux system and ammonia (0 ppm), nitrite (0 ppm), and nitrate (0 ppm) were monitored weekly (API test kits, Mars Fishcare, North America). Lighting was provided by two T5 bulbs per tank—one actinic blue and one aquablue special (ATI North America)—delivering approximately 50 PAR. The light cycle was maintained at 12 hours light and 12 hours dark (12 h L: 12 h D). The life support system included a biomedia bed for biological filtration and a UV sterilizer for disinfection.

### Experimental Set-Up

We tested the overarching hypothesis that interactions among size- and age-matched rivals will lead to the emergence of social roles accompanied by divergence in four aspects of individual phenotypes. This experimental design allowed individuals equal opportunity to attempt taking on the dominant social role, however, through continuous social interactions (or in the case of solitaries, due to the lack of it), individuals took on varying social roles as dominant and subordinate members. A total of 90 juveniles, 30 from each of three clutches, were used. Using these juveniles, we created 30 replicates with an unrelated size-matched pair housed in one breeder box (n = 30 pairs; n = 60 individuals) and a solitary individual housed in another breeder box (n = 30 individuals). At the end of the experiment, individual identities within pairs were verified using either gene expression data or microsatellite genotyping (see Supplementary Materials). This step was necessary to avoid assumptions regarding social roles and fish identity. At the beginning of the experiment, individuals within pairs were provisionally assigned a social rank based on initial body size; as size differences were negligible (often less than 0.1mm), the initially larger individual does not necessarily emerge as the dominant (P1). To correct this assumption, identities were verified at the end of the experiment using either microsatellite genotyping or SNPs called from TagSeq data, depending on whether individuals were included in the gene expression dataset (see Supplementary Materials). The larger individual in each pair was classified as pair rank 1 (P1), the smaller as pair rank 2 (P2), and solitary individuals were denoted as S.

### Body Size

To measure juvenile size, fish were photographed in a Petri dish marked with grid paper under a dissecting microscope. These photos were used to measure standard length (SL) of juveniles to 0.01 mm using ImageJ (Schneider et al., 2012). Once measured, we size-matched pairs and solitary individuals as closely as possible: average size difference between individuals within these trios was 0.66 ± 0.21mm, which is 0.08% of their average standard length.

### Feeding Behavior and Agonistic Behaviors

To measure juvenile behavior, we used cameras (Olympus Stylus Tough TG-Tracker Action camera) to record each pair and each solitary for 15 minutes, once per week, for five weeks. Note that there were no video recordings made of tanks with solitaries during the first week due to time constraints during experimental setup.

To measure feeding behavior, we introduced a divider to separate paired individuals prior to feeding, which prevented them from interacting with each other, and we introduced a divider to solitary individuals as a sham control. Then each fish was fed an identical portion of food. We scored feeding behavior — total number of bites of food taken — for 5 minutes after the first bite of food was taken, in both paired and solitary individuals. During the first two experimental weeks, P1 and P2 could not be distinguished in videos because of their similar sizes and lack of distinguishing markings. Therefore, we identified the individual who consumed more food in a given video as P-max and the one who consumed less as P-min, and we maintained this designation throughout the experiment unless members of a pair could be reliably distinguished. From week 3 onward, members of a subset of pairs could be reliably distinguished due to the emergence of clear size differences (week 3, n=5 pairs; week 4, n=17 pairs; week 5, n=19 pairs), allowing us to score food intake separately for dominant (P1) and subordinate (P2) fish.

To measure aggression and submission in paired individuals, we scored agonistic behaviors for 5 minutes after 1 minute of acclimatization (no divider present). We scored aggression based on the sum of all bites, chases, and headbutts between pairs (Wong et al., 2013, 2016). Flees and shakes between individuals were recorded to measure submissiveness between pairs (Wong et al., 2013, 2016). Because P1 and P2 could not be distinguished in videos during the first two experimental weeks, we quantified aggression as a total aggression score and submission as a total submission score for the pair, and we maintained this assignment throughout the experiment unless members of a pair could be reliably distinguished. However, aggression was typically displayed by only one member of the pair. From week 3 onward, members of a subset of pairs could be reliably distinguished due to the emergence of clear size differences (week 3, n = 5 pairs; week 4, n = 17 pairs; week 5, n = 19 pairs), allowing us to score aggression and submission separately for dominant (P1) and subordinate (P2) fish. Aggression and submission were not scored for solitary individuals.

### RNA Isolation and Sequencing

Following the experiment described above, all fish were preserved for gene expression profiling. At the time of euthanasia, all fish had not been fed for over 14 hours, ensuring consistency for downstream analysis. Before preservation, all fish (n=90) were individually cold-anesthetized according to protocols previously used in the lab (Maytin et al., 2018) after which whole fish tissue samples were immediately preserved in 200 proof ethanol. Following preservation, 15 replicates were chosen for subsequent gene expression profiling based on preliminary results indicating that growth varied consistently with social position. To maximize our ability to detect underlying gene expression patterns contributing to the expected growth signature (P1 > P2 = S), 15 replicates that exhibited the greatest size difference between P1 and P2 were selected for gene expression profiling. We also controlled, to the degree possible, for genetic effects by choosing an equal number of fish from each clutch across social positions (P1, P2, S). Fish from these 15 replicates (n=45 samples; 15 paired individuals and their corresponding solitaries) were individually homogenized to allow equal RNA extraction from all tissues. To initially explore overall gene expression patterns associated with coordinated changes during the emergence of social roles, whole-body RNA-seq was performed to keep both sequencing cost and sample requirements manageable. We acknowledge the limitations of this approach and consider this study a hypothesis generating tool that lays the foundation for future studies.

Tissue samples were cut into smaller pieces with a razor blade and RNA was isolated using an established phenol-chloroform DNA/RNA isolation protocol, with the modification of not adding RNAse (Davies et al., 2013). Samples were eluted in 35µL of molecular biology grade water and DNAsed using RNA Clean & Concentrator-5 kit (Zymo Research, no. R1015) according to manufacturer’s protocol. RNA concentrations were determined using a DeNovix DS-11+ Spectrophotometer (DeNovix Inc., Wilmington, DE, USA) and RNA integrity was visualized on a 1% agarose gel. Next, all samples were individually normalized to 100ng/25µL and sent to Genomic Sequencing and Analysis Facility at the University of Austin, Texas, where TagSeq libraries (Meyer et al., 2011) were prepared and sequenced (single end 100bp) on a NovaSeq 6000. RNA sequencing and downstream analyses were conducted separately for each of the 45 samples (15 P1, 15 P2, and 15 S).

### Raw Sequence Processing and Read Mapping

A total of 366 million raw reads were generated with average raw read counts of 8.2 million (range: 5.4 to 11.3 million per sample; Supplementary Table S1). Raw reads were trimmed with *Fastx_toolkit* (Hannon, 2010), which removed 5’-Illumina leader sequences and poly(A)+ tails. Sequences <20bp in length with <90% of bases having quality cutoff scores <20 were also trimmed. In addition, because degenerate bases were incorporated during cDNA synthesis, PCR duplicates were removed from all libraries. After quality filtering, an average of 3.2 million reads per sample remained (ranging from 2.3 to 4.4 million per sample; Supplementary Table S1), which were then mapped to the *Amphiprion percula* genome (Lehmann et al., 2019) using *STAR* (Dobin et al., 2013). Mapped bam files were used to generate per gene read counts across samples using *featureCounts*. The read count file was then imported into R version 4.3.1 (R Core Team, 2021), size factors were estimated for each sample using the median ratio method in DESeq2 (Love et al., 2014) to account for differences in sequencing depth across samples. No outlier samples were identified, and all samples (n=45) were retained for downstream analyses. Genes with a mean raw count <10 were then removed, retaining 24,840 genes for downstream analyses. To correctly identify individuals within pairs and ensure they are correctly assigned to clutches, we called single-nucleotide polymorphisms (SNPs) from TagSeq data (see Supplementary Materials), prior to downstream analysis.

### Statistical Analyses

From the total of 30 replicates, 3 replicates lost fish throughout the five-week experiment. All three losses were within pairs. These three replicates were removed from all downstream analyses, leaving a total of 27 replicates (N=54 paired individuals; N=27 solitary individuals). Given that all phenotypes are entirely social context-dependent with no intrinsic baseline, in this species, we arbitrarily chose P1 (dominant) as the statistical reference group for all downstream analyses. To test our four hypotheses, we ran four general linear mixed models (GLMM) using the *LmerTest* package (Kuznetsova et al., 2017) in R. We considered size as the dependent variable, with social position of individual (P1, P2, or S), time (t1 or t5), and their interaction as the independent variables. We considered feeding behavior as the dependent variable, with social position of individual (P-max, P-min, or S), time (t1, t2, t3, t4, or t5), and their interaction as the independent variables. In addition, for the subset of replicates where dominant (P1) and subordinate (P2) individuals could be reliably distinguished (weeks 3–5), we ran a complementary GLMM using known social roles (P1, P2, S), time (t1, t2, t3, t4, or t5), and their interaction as the independent variables. Furthermore, we considered agonistic behavior (aggression or submission) between individuals within pairs as the dependent variables, with time (t1, t2, t3, t4, or t5) as the independent variable. For the subset of pairs where P1 and P2 could be distinguished (weeks 3–5), we also ran complementary GLMMs using known social roles (P1, P2) as the independent variables. To account for the lack of independence among individuals from the same replicate, all our models included replicate ID as a random effect. Model assumptions were assessed by visual inspection of residuals and fitted values; residuals were approximately normally distributed and showed no evidence of heteroscedasticity, thus no data transformations were required. In all cases, the best fit model was determined by carrying out a backward model selection (*step* function in *LmerTest* package (Kuznetsova et al., 2017). Post hoc pairwise contrasts using *emmeans* (Searle et al., 1980) were used to determine where differences exist between categories.

### Global Gene Expression Patterns

All gene expression (GE) analyses were carried out in R (version 4.3.1). To test for differentially expressed genes (DEGs) across the social positions (P1, P2, and S) DESeq2 (Love et al., 2014) with the model: design = ∼ genotype + social position was run. Count data were normalized and independent pairwise contrasts across 3 social positions were computed. Genes identified as differentially expressed were corrected for multiple testing using false discovery rate (FDR: adjusted p < 0.05). Annotated DEGs of up- and down-regulated genes for all three pairwise comparisons were visualized using volcano plots. To test for overall GE patterns between social positions and genotypes, data were rlog-normalized and the effect of genotype and social position were compared through a Principal Component Analysis (PCA), followed by PERMANOVA analysis using Euclidean distances in the vegan package (Oksanen et al., 2017). To assess whether the strength of clustering by social position differed depending on the genes included, additional PCAs were performed on four gene subsets: (1) all genes, (2) the top 50%, (3) the top 10%, and (4) the top 5% of the most variable genes. For each subset, the first three principal components (PC1 vs PC2, PC1 vs PC3, PC2 vs PC3) were visualized to identify axes capturing variation associated with social position. To test for overall gene expression differences between social position and genotype, separate PERMANOVAs were performed for each PCA using the *adonis2* in the vegan package (Oksanen et al., 2017).

### Gene Co-Expression Networks Associate Expression Patterns with Phenotypic Traits

To test whether gene expression patterns are correlated with observed phenotypic changes in growth and appetite across social positions, a Weighted Gene Correlation Network Analysis (WGCNA) was conducted on the rlog-transformed GE dataset (Langfelder & Horvath, 2008). WGCNA identifies groups of co-expressed genes (modules) and correlates eigengene expression of each module with phenotypes of interest to determine modules of genes whose expression correlates with trait variation. A signed network was constructed using a soft-threshold power of 6 with both the network and Topological Overlap Matrix (TOM) set to signed. Modules were identified with a minimum module size of 600, and module merging threshold was set to 0. For modules whose eigengene expression correlated with phenotypes of interest, gene ontology (GO) enrichment analysis was subsequently performed using Fisher’s Exact Tests (presence/absence in a module) to identify over-represented pathways within each module across GO divisions of Biological Processes (BP), Molecular Functions (MF), and Cellular Components (CC) (Wright et al., 2015). Genes associated with specific GO terms were explored and significant DEGs (FDR: p ≤ 0.05) were extracted and visualized with *ComplexHeatmaps* (Gu et al., 2016). In the heatmap, samples (columns) were *a priori* grouped by social position (P1, P2, S) and subsequently ordered within each group by gene expression similarity, reflected by the column dendrogram.

Additionally, we tested the hypothesis that size ratio of pairs (P2 SL / P1 SL), which is indicative of the amount of conflict remaining (Wong et al., 2016), would predict variation in gene expression. This was assessed by visual inspection of the same heatmap with size ratio annotation using *ComplexHeatmaps* (Gu et al., 2016), together with a PCA of the expression of these genes of paired individuals (P1, P2) only. We also performed a PERMANOVA analysis using the same row Z-score values (*adonis2* function in vegan package: Oksanen et al., 2017) accounting for social position and genetic background (clutch ID), using restricted permutations to account for the non-independence of P1 and P2 within pairs was carried out. Solitary individuals were excluded from this analysis because they lack size ratio data. We found that size ratio between pairs was not associated with gene expression variation within social position (see Supplementary Materials; Supplementary Fig. S8, S10A).

To further investigate GE differences, we explored genes associated with growth (including thyroid signaling), appetite regulation, metabolism, and stress (corticoid) pathways across social positions. Specifically, we focused on genes associated with these pathways previously identified in the sister species, *Amphiprion ocellaris* (Herrera et al., 2025). Genes identified in *A. ocellaris* were used to retrieve *A. percula* orthologs, using a reciprocal BLAST search (BLAST+ version 2.11.0) approach (Camacho et al., 2009). Both BLAST searches were performed with an e-value threshold of 0.0001 to ensure high ortholog confidence. Only gene pairs that were the best match in both blast directions were considered as orthologs for downstream analysis. A complete list of retrieved *A. percula* gene IDs were then filtered against the whole-body GE dataset, and pathways containing significant genes associated with social position were reported (Supplementary Table S2; appetite and metabolic genes shown). Of these, significant pathway DEGs (FDR p ≤ 0.01) were visualized using *ComplexHeatmaps* (Gu et al., 2016) while taking into account genotype and social position. In the heatmap, samples (columns) and genes (rows) are grouped *a priori* by social position (P1, P2, S) and pathway (APT, TCA, GLY), respectively, with dendrograms reflecting expression similarity within each group. For the candidate gene heatmap, we repeated the PCA and PERMANOVA exploration and associated heatmap visualization for pairs only (as described above), to test whether final size ratio (P2 SL / P1 SL) predicted gene expression variation within social positions. We found that size ratio between pairs was not associated with gene expression variation within social position (see Supplementary Materials; Supplementary Fig. S9, S10B).

## Acknowledgements

We are particularly grateful to the three reviewers, Karen Warkentin, Shannon Mary Fisher, Paige Becker, and Zoé Chamot, for their constructive feedback on the manuscript. We also thank Natalia Karadimitriou for her invaluable assistance during larval rearing and experimental setup. This study forms part of the Ph.D. dissertation of Lili Vizer (Boston University).

## Ethics

All applicable national and/or institutional guidelines for the care and use of animals were followed. All procedures with live animals were reviewed and approved by the Boson University IACUC (protocol number PROTO201800525).

## Data Accessibility

Raw FASTQ files for all gene expression samples are publicly accessible through the NCBI Sequence Read Archive (SRA) under BioProject accession number **PRJNA1280023**. All data, along with the code used for analyses and figure generation, are available on GitHub at: https://github.com/lillentyu/A-percula-Adaption-Of-Social-Roles.

## Authors’ contributions

L.F.V.: conceptualization, data curation, formal analysis, funding acquisition, investigation, methodology, project administration, visualization, writing—original draft, writing—review and editing; D.A.: data curation, investigation, methodology, writing—review and editing; C.B.: methodology, writing—review and editing; M.H: methodology, writing—review and editing; A.H.: methodology, writing—review and editing; K.T.: methodology, writing—review and editing; S.M.B.: data curation, methodology, writing—review and editing; V.L.: methodology; supervision, writing—review and editing; S.W.D.: methodology; visualization, supervision, writing—review and editing; P.M.B.: conceptualization, funding acquisition, investigation, methodology, project administration, supervision, writing—review and editing.

## Declaration of AI use

The artificial intelligence tool ChatGPT (OpenAI, version GPT-4) was used to support R script development, particularly for data management, analysis, and visualization. Prompts included requests such as: “Identify errors or issues in the following code based on this data frame structure: [description of code and data], and suggest possible alternatives or solutions,” and “Explain and resolve this error message: [error message and associated code].” ChatGPT assisted by identifying and explaining coding issues, offering corrected code, and suggesting alternative approaches where appropriate. All suggestions provided by ChatGPT were reviewed by an expert before implementation.

## Supplementary materials

## This file includes

Methods of Microsatellite and Single Nucleotide Polymorphisms (SNP) calling and Supplementary Figures S1, S2

Stand-alone Supplementary Tables S1, S2, S3 and S4

Stand-alone Supplementary Figures S3, S4, S5, S6, and S7

Gene Expression Analysis as a Function of Size Ratio Supplementary Figures S8, S9, S10

Color Dataset Analysis and Supplementary Figure S11

## Microsatellite Genotyping and Single Nucleotide Polymorphism (SNP) calling from TagSeq

For the 36 samples which were not part of the gene expression dataset, multiplex microsatellite genotyping was carried out to ensure that individuals within pairs are correctly identified and clutch assigned. This step was necessary to avoid assumptions regarding social roles and fish identity, and the microsatellite method allowed for a reliable and cost-effective solution as a panel with established markers was already available from previous study (Rueger et al., 2025). At the beginning of the experiment, individuals within pairs were provisionally assigned a social rank based on initial body size; as size differences were negligible (often less than 0.1 mm), the initially larger individual does not necessarily emerge as the dominant (P1). To correct this assumption, identities were verified at the end of the experiment. Fin-clips were taken from preserved samples and DNA extracted using HotShot (Truett et al., 2000). 75 µL of alkali lysis buffer was added to fin-clips and incubated at 90°C for 30 minutes, followed by cooling and addition of 75 µL of neutralizing buffer. These samples were then sent for multiplex PCR and library preparation to the Environmental DNA and Genomics Core Facility at Cornell University, and paired end Illumina sequencing at the Genomics facility of the Cornell Bioresource Center according to previously established protocols (D’Aloia et al., 2017). Haplotype data of 60 *A. percula* specific loci for the 36 individuals were compiled into a matrix format. Due to robustness of initial reconstruction, no individuals or SNPs were required to be filtered. Genetic distances between individuals were computed using the *dist.gene* function from the *ape* package in R (Paradis et al., 2004), applying the “pairwise” method with pairwise deletion enabled to account for missing data. Hierarchical clustering was then performed using the *hclust Ward.D2* method (Müllner, 2013), and a dendrogram was generated to visualize genetic relatedness among individuals and to assess whether individuals had been correctly assigned to clutches (Supplementary Fig. S1A). Pairs assigned incorrectly were then corrected and used for downstream analysis (Supplementary Fig. S1B).

**Supplementary Figure S1:**
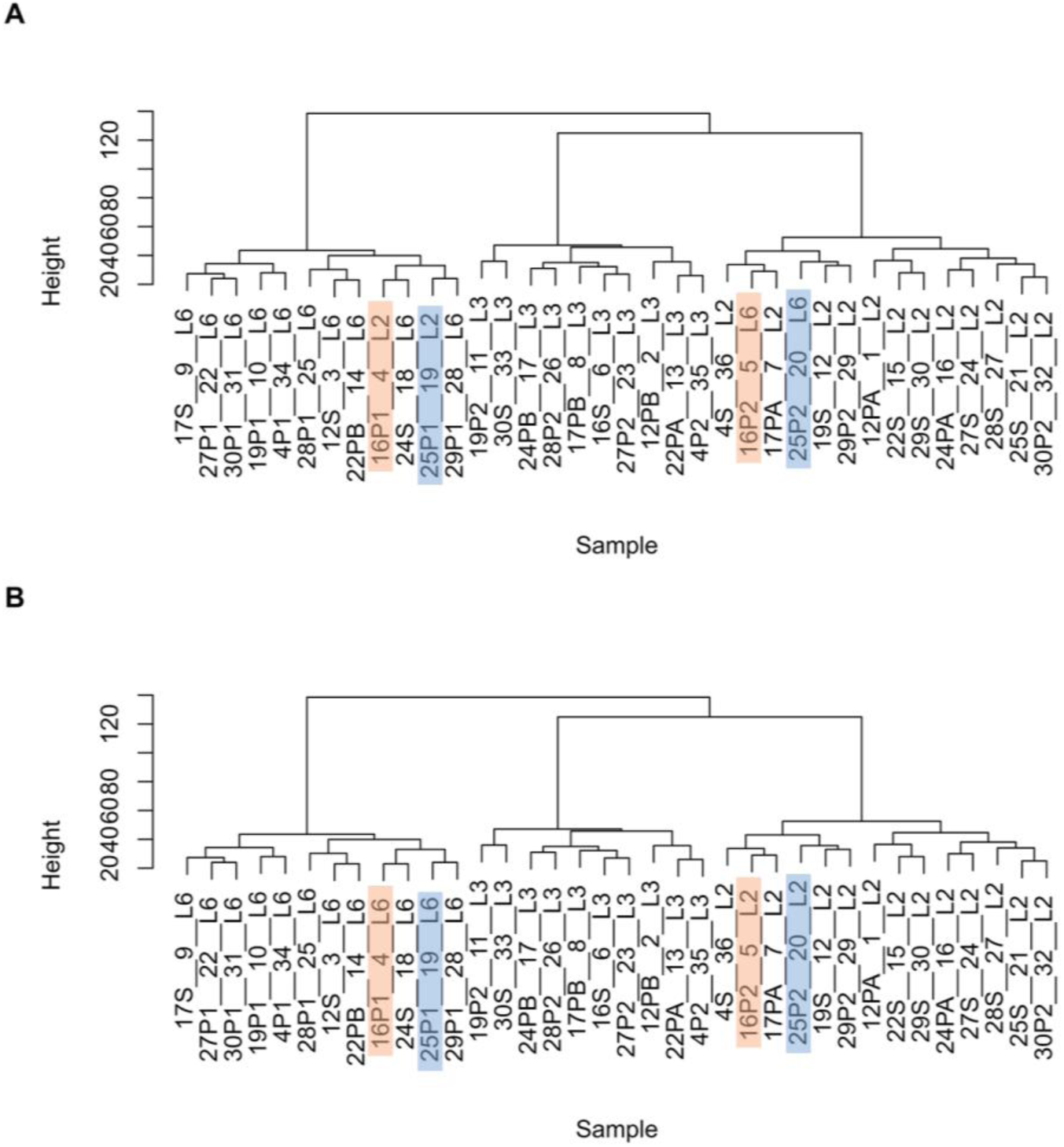
Clutch assignment validation via hierarchical clustering of microsatellite genotypes. The dendrogram illustrates genetic relatedness among 36 individuals from three clutches, based on microsatellite genotypes (60 loci) processed using HotShot genotyping. **A)** Initial assignment of individuals to their presumed clutches, with pairs incorrectly assigned highlighted in blue and orange. Branch lengths reflect genetic divergence; tightly clustered branches indicate sibling relationships within clutches. **B)** Corrected clustering after reassignment of clutches for mismatched pairs.

Similarly, for the remaining 45 individuals, we applied a necessary correction step to avoid assumptions regarding social role and fish identity. For these 45 individuals, we called SNPs from our TagSeq data to assign clutch identity. Quality filtered and trimmed reads were mapped against the *A. percula* transcriptome (Maytin et al., 2018) using Bowtie2.2 (Langmead & Salzberg, 2012). Generated SAM files were converted to BAM format using *samtools* (Li et al., 2009). *ANGSD* (Korneliussen et al., 2014) then calculated pairwise identity by state (IBS) matrices, which were used as input for *hclust* (Müllner, 2013) to visualize relatedness among the 45 individuals (Supplementary Fig. S2A). All individuals within pairs were then assigned to the correct clutch (Supplementary Fig. S2B), which was used for downstream analyses.

**Supplementary Figure S2:**
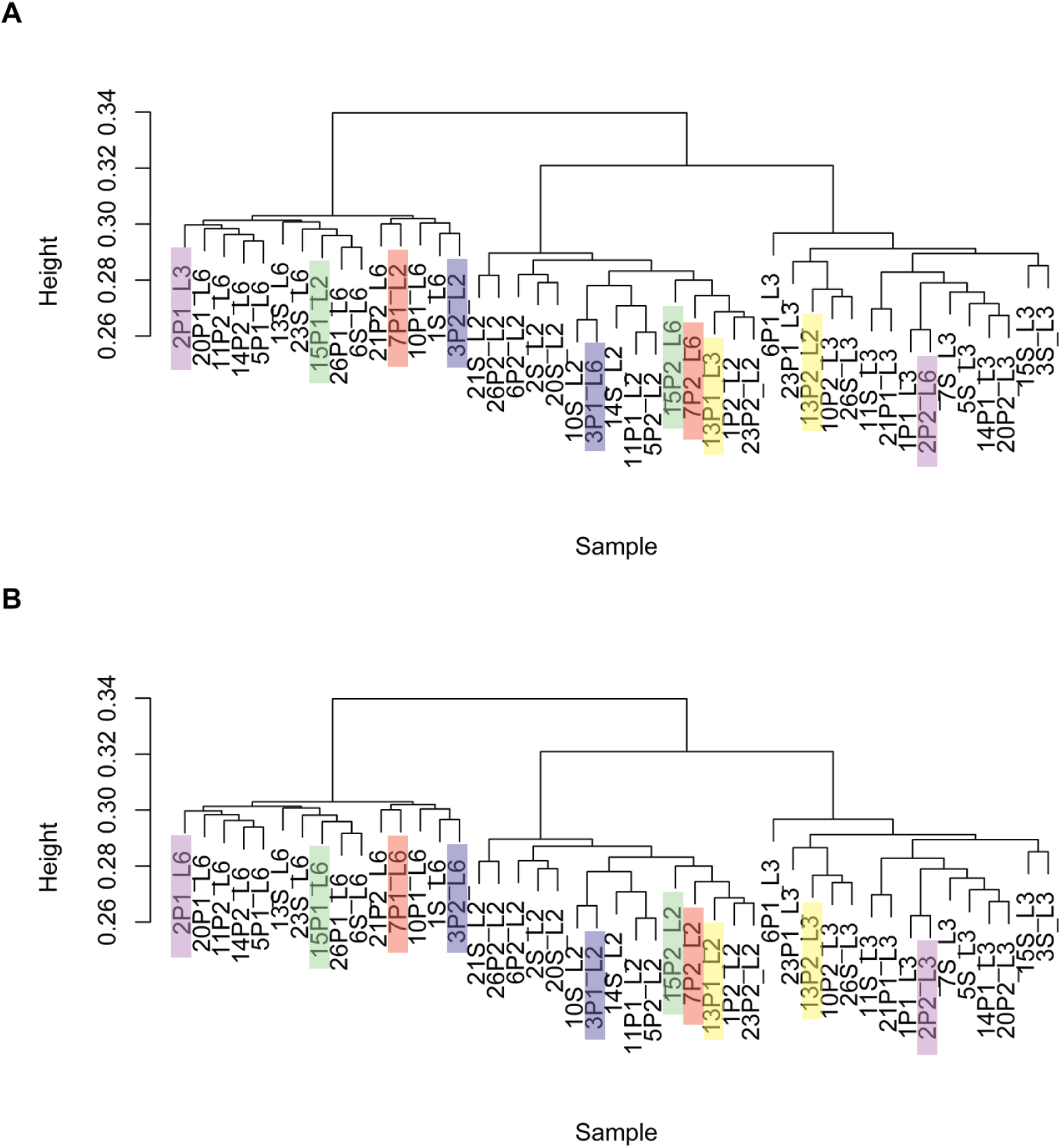
Clutch assignment validation using TagSeq-derived SNPs and hierarchical clustering. The dendrogram illustrates genetic relatedness among 45 individuals from three clutches, based on SNPs from TagSeq data. **A)** Initial assignment of individuals to their presumed clutches, with pairs incorrectly assigned highlighted in colors. Branch lengths reflect genetic divergence; tightly clustered branches indicate sibling relationships within clutches. **B)** Corrected clustering after reassignment of clutches for mismatched pairs.

## Stand-alone Supplementary Tables

**Supplementary Table S1:** Mapped read count data. *A. percula* replicate, social position (P1 – pair rank 1; P2 – pair rank 2; S – solitary) and clutch ID. Read counts of raw, filtered and trimmed sequences, and number of uniquely mapped reads against *A. percula* genome using STAR, as well as the corresponding mapping efficiency.

| Sample ID | Social Position | Clutch ID | Raw Read Counts | Trimmed Read Counts | Uniquely Mapped Reads | Mapping Efficiency (%) |
| --- | --- | --- | --- | --- | --- | --- |
| 1 | P1 | L3 | 5986911 | 2737307 | 2352774 | 85.952 |
| 1 | P2 | L2 | 6803145 | 2954216 | 2554648 | 86.475 |
| 1 | S | L6 | 5837482 | 2648669 | 2294955 | 86.646 |
| 2 | P1 | L6 | 6217888 | 2787974 | 2394766 | 85.896 |
| 2 | P2 | L3 | 7010907 | 3100482 | 2669479 | 86.099 |
| 2 | S | L2 | 8019177 | 3585504 | 3072414 | 85.690 |
| 3 | P1 | L2 | 8383004 | 3518423 | 2955511 | 84.001 |
| 3 | P2 | L6 | 7386810 | 3360910 | 2862178 | 85.161 |
| 3 | S | L3 | 7249814 | 3348687 | 2858570 | 85.364 |
| 5 | P1 | L6 | 7202343 | 3143829 | 2699683 | 85.872 |
| 5 | P2 | L2 | 6390202 | 2808877 | 2416787 | 86.041 |
| 5 | S | L3 | 5400980 | 2350517 | 2050627 | 87.242 |
| 6 | P1 | L3 | 6649594 | 2507798 | 2160966 | 86.170 |
| 6 | P2 | L2 | 7091099 | 3238004 | 2789147 | 86.138 |
| 6 | S | L6 | 7237912 | 3074694 | 2646890 | 86.086 |
| 7 | P1 | L6 | 7204499 | 2999504 | 2547277 | 84.923 |
| 7 | P2 | L2 | 6970070 | 2899158 | 2483166 | 85.651 |
| 7 | S | L3 | 6461090 | 2879935 | 2473878 | 85.900 |
| 10 | P1 | L6 | 6168327 | 2471974 | 2128137 | 86.091 |
| 10 | P2 | L3 | 6778638 | 2834810 | 2446430 | 86.300 |
| 10 | S | L2 | 6303813 | 2753496 | 2356609 | 85.586 |
| 11 | P1 | L2 | 7700902 | 2871152 | 2433468 | 84.756 |
| 11 | P2 | L6 | 10410675 | 3926127 | 3378190 | 86.044 |
| 11 | S | L3 | 8457694 | 3369689 | 2904659 | 86.200 |
| 13 | P1 | L2 | 8747232 | 2756847 | 2388352 | 86.633 |
| 13 | P2 | L3 | 11355428 | 3983796 | 3392123 | 85.148 |
| 13 | S | L6 | 8493446 | 3498055 | 2976781 | 85.098 |
| 14 | P1 | L3 | 9649613 | 3377460 | 2892559 | 85.643 |
| 14 | P2 | L6 | 11096303 | 3812633 | 3307234 | 86.744 |
| 14 | S | L2 | 7458006 | 2925835 | 2543736 | 86.941 |
| 15 | P1 | L6 | 8315175 | 2770012 | 2448710 | 88.401 |
| 15 | P2 | L2 | 7809486 | 3122052 | 2657505 | 85.120 |
| 15 | S | L3 | 11188485 | 4419431 | 3841559 | 86.924 |
| 20 | P1 | L6 | 10293955 | 3756124 | 3194578 | 85.050 |
| 20 | P2 | L3 | 9793618 | 3759257 | 3248705 | 86.419 |
| 20 | S | L2 | 10601642 | 3926835 | 3360769 | 85.585 |
| 21 | P1 | L3 | 9434952 | 3630988 | 3124380 | 86.048 |
| 21 | P2 | L6 | 9464753 | 3554760 | 3070925 | 86.389 |
| 21 | S | L2 | 10612954 | 3949458 | 3418245 | 86.550 |
| 23 | P1 | L3 | 6565976 | 2610543 | 2201455 | 84.329 |
| 23 | P2 | L2 | 10032443 | 4035965 | 3453739 | 85.574 |
| 23 | S | L6 | 10407184 | 3981383 | 3457368 | 86.838 |
| 26 | P1 | L6 | 7843938 | 2824540 | 2461006 | 87.129 |
| 26 | P2 | L2 | 7756092 | 3046595 | 2636393 | 86.536 |
| 26 | S | L3 | 10632681 | 4163850 | 3607742 | 86.644 |

**Supplementary Table S2:**
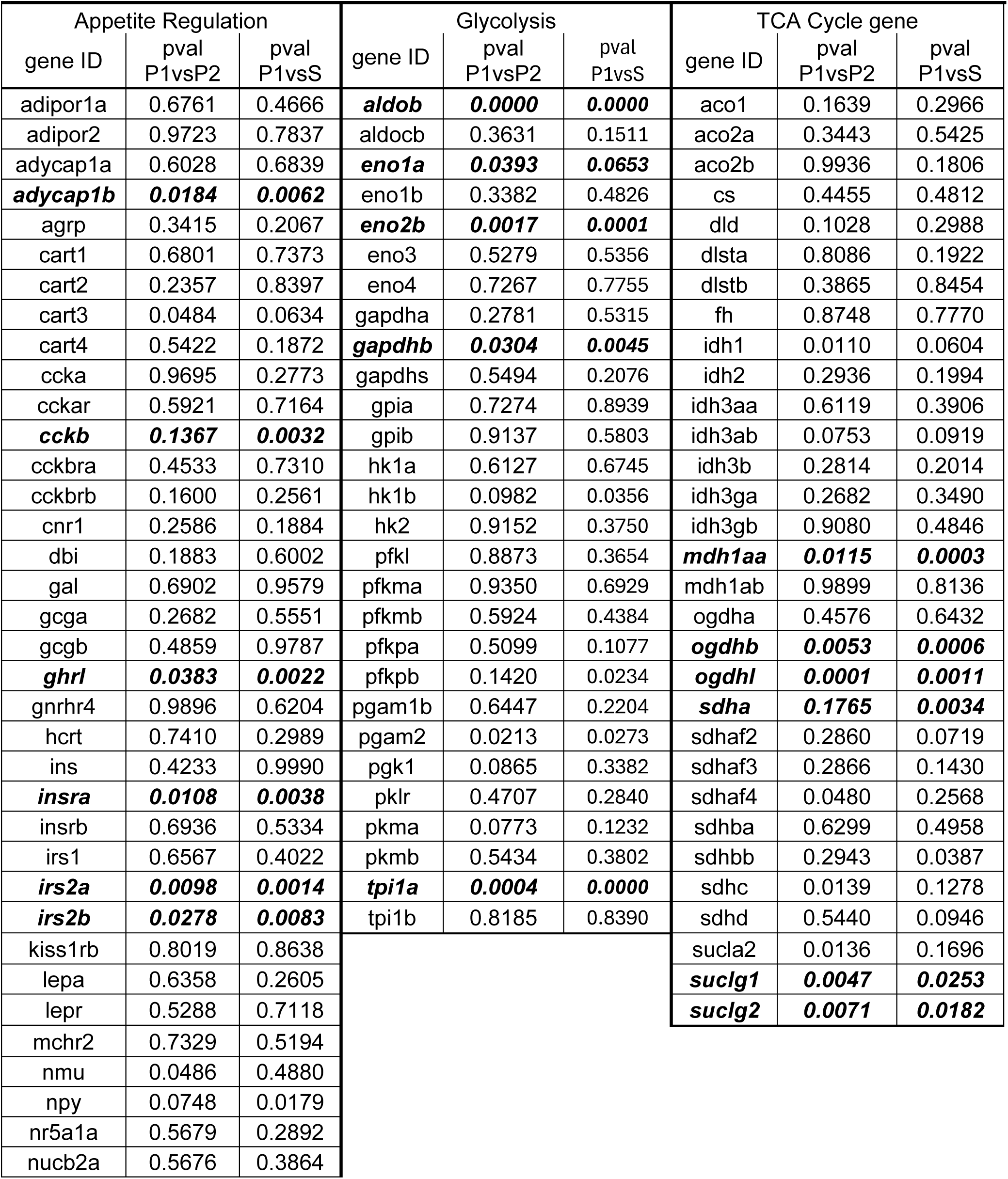
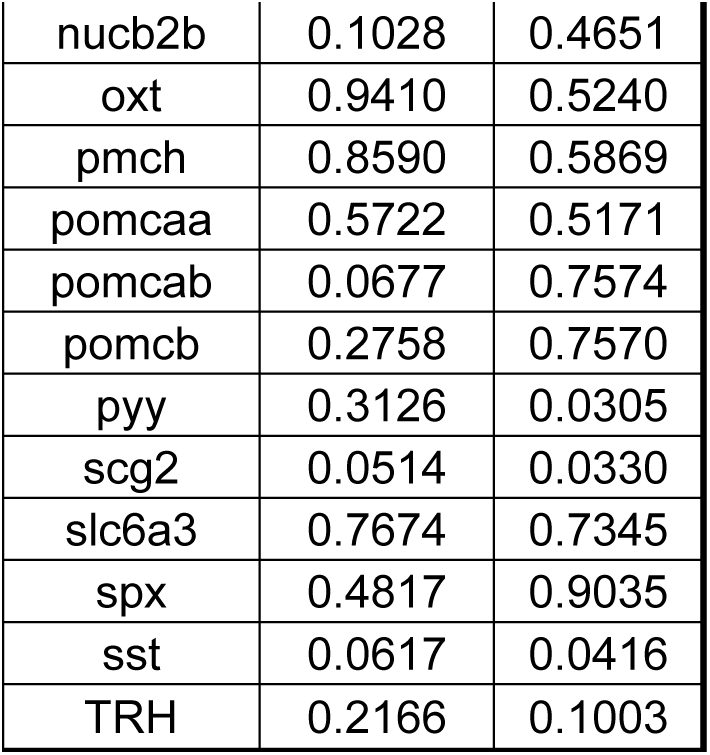
Complete list of *Amphiprion percula* genes associated with appetite regulation, glycolysis, and the TCA cycle, retrieved based on orthologs with *A. ocellaris* genes (Herrera et al., 2025). Orthologs were identified using a reciprocal BLAST approach, and genes present in the whole-body gene expression dataset were retained. The table includes each gene and its associated p-values from pairwise contrasts between social positions (P1 – pair 1; P2 – pair 2; S – solitary). Genes with p-value ≤ 0.01, which were visualized in the heatmap shown in the main text (Fig. 6), are indicated in ***bold italics***.

**Supplementary Table S3:** Post-hoc pairwise contrasts of total aggression (upper triangle) and total submission (lower triangle), displaying p-values of all comparisons. Significant differences are denoted with *.

|  | *Week 1* | *Week 2* | *Week 3* | *Week 4* | *Week 5* |
| --- | --- | --- | --- | --- | --- |
| *Week 1* |  | 0.688 | 0.026* | < 0.001* | < 0.001* |
| *Week 2* | 0.285 |  | 0.686 | < 0.001* | < 0.001* |
| *Week 3* | < 0.001* | 0.1469 |  | 0.028* | 0.002* |
| *Week 4* | < 0.001* | 0.002* | 0.4039 |  | 0.937 |
| *Week 5* | < 0.001* | < 0.001* | 0.1477 | 0.9845 |  |

**Supplementary Table S4:** List of significantly differentially expressed genes shared between the pairwise comparisons of P1 vs P2 and P1 vs S (main text Fig. 3B). A total of 25 genes were consistently downregulated in P1 (relative to both P2 and S), and 84 genes were consistently upregulated in P1. Gene names are shown where available; otherwise, Ensembl gene IDs are provided.

| shared downregulated genes | shared upregulated genes |  |  |  |
| --- | --- | --- | --- | --- |
| c1qtnf5 | BRSK2 | IGFBP1 | SHMT2 | ENSAPEG00000005722 |
| COL10A1 | aldh6a1 | IQSEC3 | slc13a5a | ENSAPEG00000006596 |
| col1a1a | aldh8a1 | LAMP5 | slc16a9a | ENSAPEG00000015523 |
| col1a2 | aldob | LECT2 | slc6a1b | ENSAPEG00000015722 |
| dpt | ambp | LRTM2 | SLC7A5 | ENSAPEG00000006747 |
| hspa1b | ATCAY | mapk8a | sybu | ENSAPEG00000008011 |
| loxa | C8A | minpp1a | syt5b | ENSAPEG00000012490 |
| MRC2 | cabp1b | MPP4 | ucp1 | ENSAPEG00000016217 |
| mustn1b | cacna1da | OGDHL | ENSAPEG00000000371 | ENSAPEG00000020211 |
| mylpfa | camk1gb | oxct1a | ENSAPEG00000003493 |  |
| myo1ca | CHIA | pax10 | ENSAPEG00000004162 |  |
| PRKAB2 | cp | pax6b | ENSAPEG00000011721 |  |
| pvalb4 | defbl2 | pck2 | ENSAPEG00000011919 |  |
| runx3 | diabloa | PCYT2 | ENSAPEG00000013967 |  |
| s100a1 | fam219aa | pde6ha | ENSAPEG00000018654 |  |
| tcf3a | ftcd | PLPPR1 | ENSAPEG00000021138 |  |
| tcf7 | GABRG2 | ppa1a | ENSAPEG00000022463 |  |
| TNNI2 | gatd3l | prph2lb | ENSAPEG00000024362 |  |
| VAT1 | glrbb | rbp2b | ENSAPEG00000006913 |  |
| ENSAPEG00000006693 | glud1b | rcvrna | ENSAPEG00000016229 |  |
| ENSAPEG00000008023 | got1 | rnf34b | ENSAPEG00000022341 |  |
| ENSAPEG00000008236 | HABP2 | saga | ENSAPEG00000023250 |  |
| ENSAPEG00000010334 | hlfa | SCG2 | ENSAPEG00000024133 |  |
| ENSAPEG00000021032 | hpda | SERPINA4 | ENSAPEG00000024384 |  |
| ENSAPEG00000023977 | hsd11b1la | SEZ6L | ENSAPEG00000001601 |  |

## Stand-alone Supplementary Figures

**Supplementary Figure S3:**
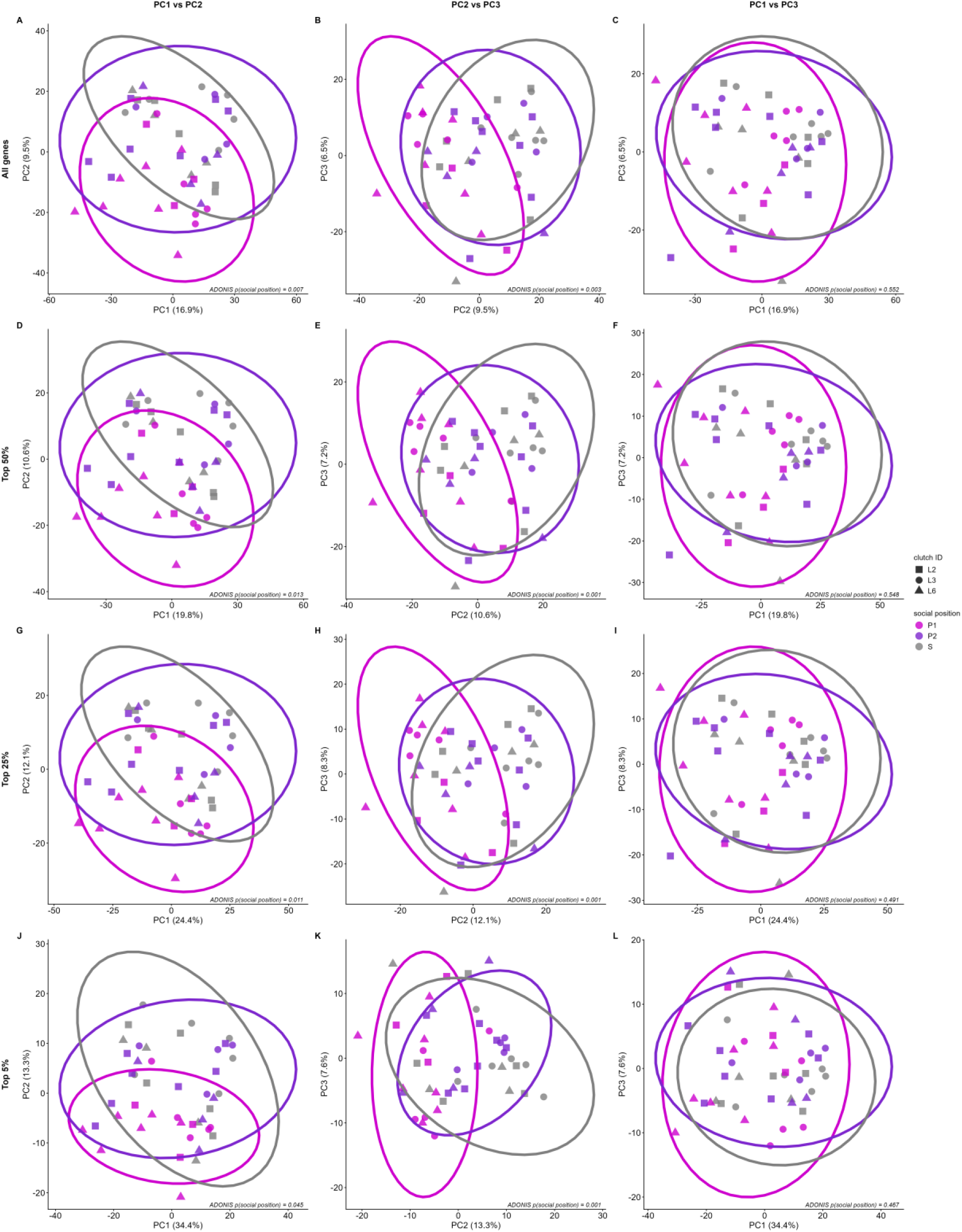
Principal component analysis (PCA) of gene expression across gene subsets and PC axis combinations. Each panel shows a PCA scatter plot of individual samples (n = 45) colored by social position (P1, P2, S), and shaped by genetic background (clutch ID). Rows represent gene subsets used for PCA: all genes **(A–C)**, top 50% **(D–F)**, top 25% **(G–I)**, and top 5% most variable genes **(J–L)**. Columns represent PC axis combinations: PC1 vs PC2 (left), PC2 vs PC3 (center), and PC1 vs PC3 (right). Ellipses show 95% confidence intervals estimated from a multivariate t-distribution for each social position. PERMANOVA p-values indicate the effect of social position on sample distribution, accounting for the genetic background of individuals (clutch ID). Panel B corresponds to Figure 3 in the main text.

**Supplementary Figure S4:**
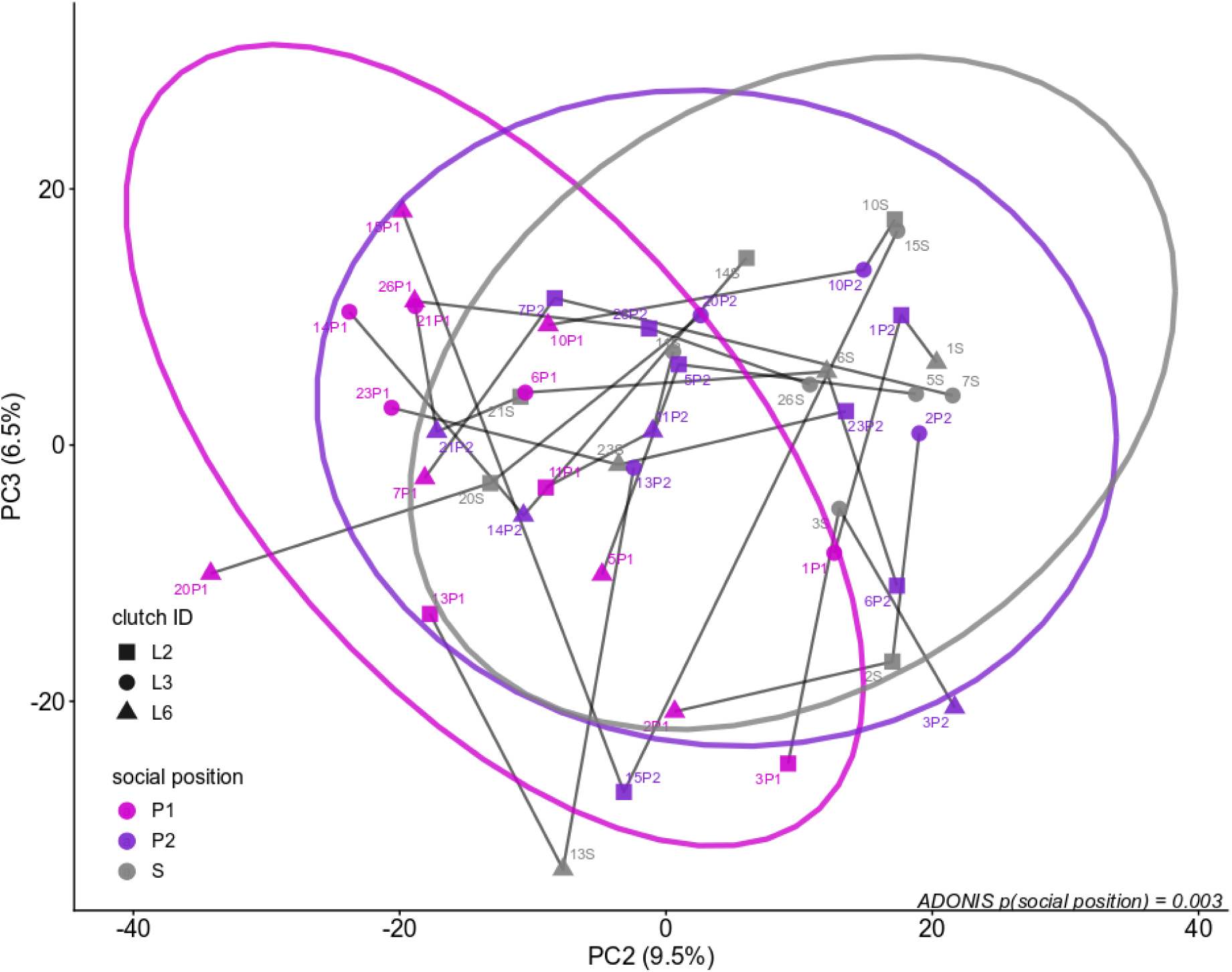
Overall gene expression profiles showed no significant differences between P2 and S, compared to P1 individuals. Principal Component Analysis (PCA) of all genes across social positions, accounting for clutch ID (genetic background), with PERMANOVA results showing a significant effect of social position. Samples are labeled, and replicates sharing the same numeric ID are connected by lines. Ellipses show the estimated confidence intervals from a 95% multivariate t-distribution.

**Supplementary Figure S5:**
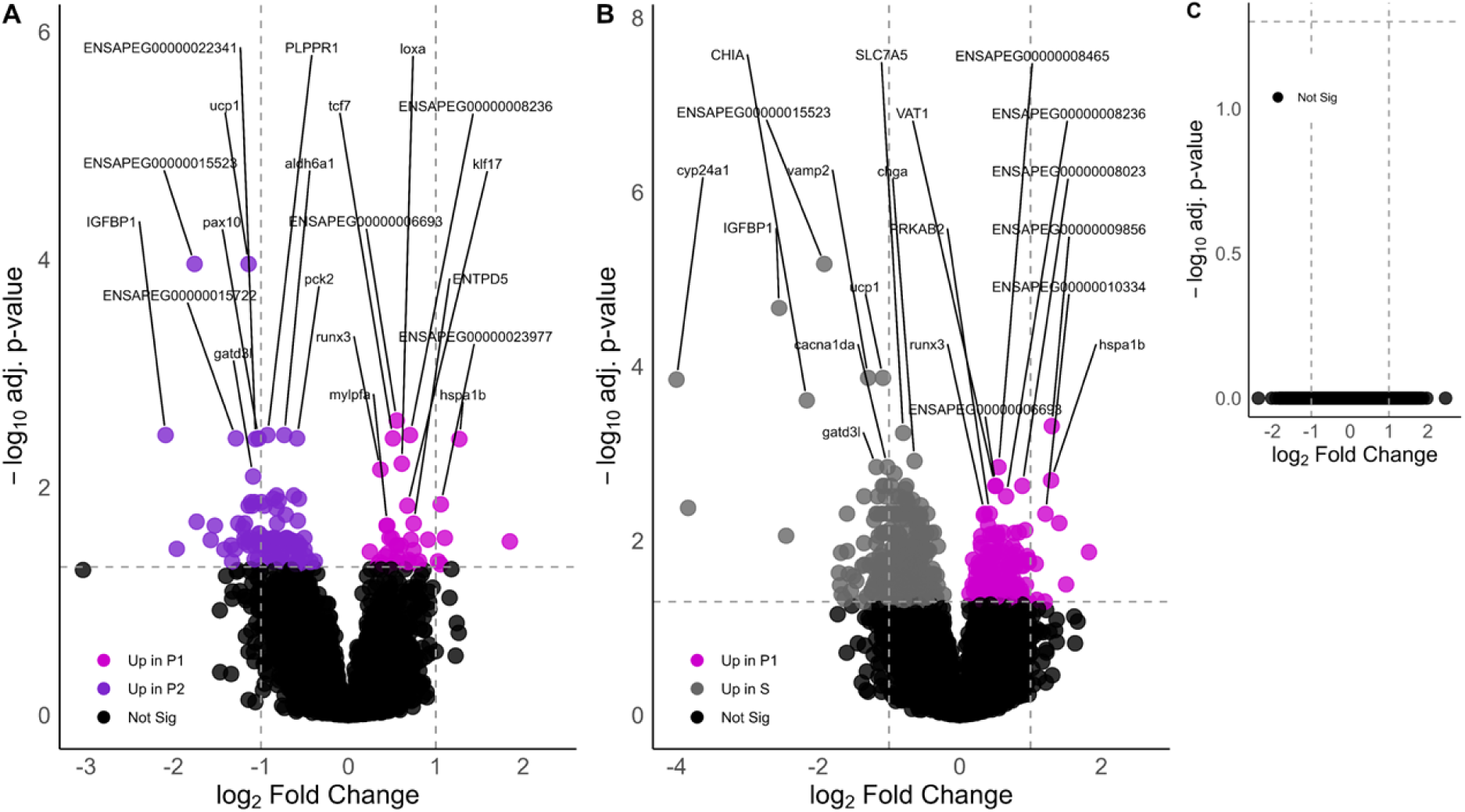
Volcano plot of differential gene expression between pair-wise comparisons of social positions (P1: dominant, P2: subordinate, S: solitary) of *A. percula*. Pairwise comparisons of differentially expressed genes (DEGs) between **A)** P1 and P2, **B)** P1 and S, and **C)** P2 and S individuals. Significant DEGs (adjusted p < 0.05) are colored according to the social position in which they are upregulated. The top 10 most significant DEGs in each pairwise comparison are labeled with gene name or Ensembl gene ID.

**Supplementary Figure S6:**
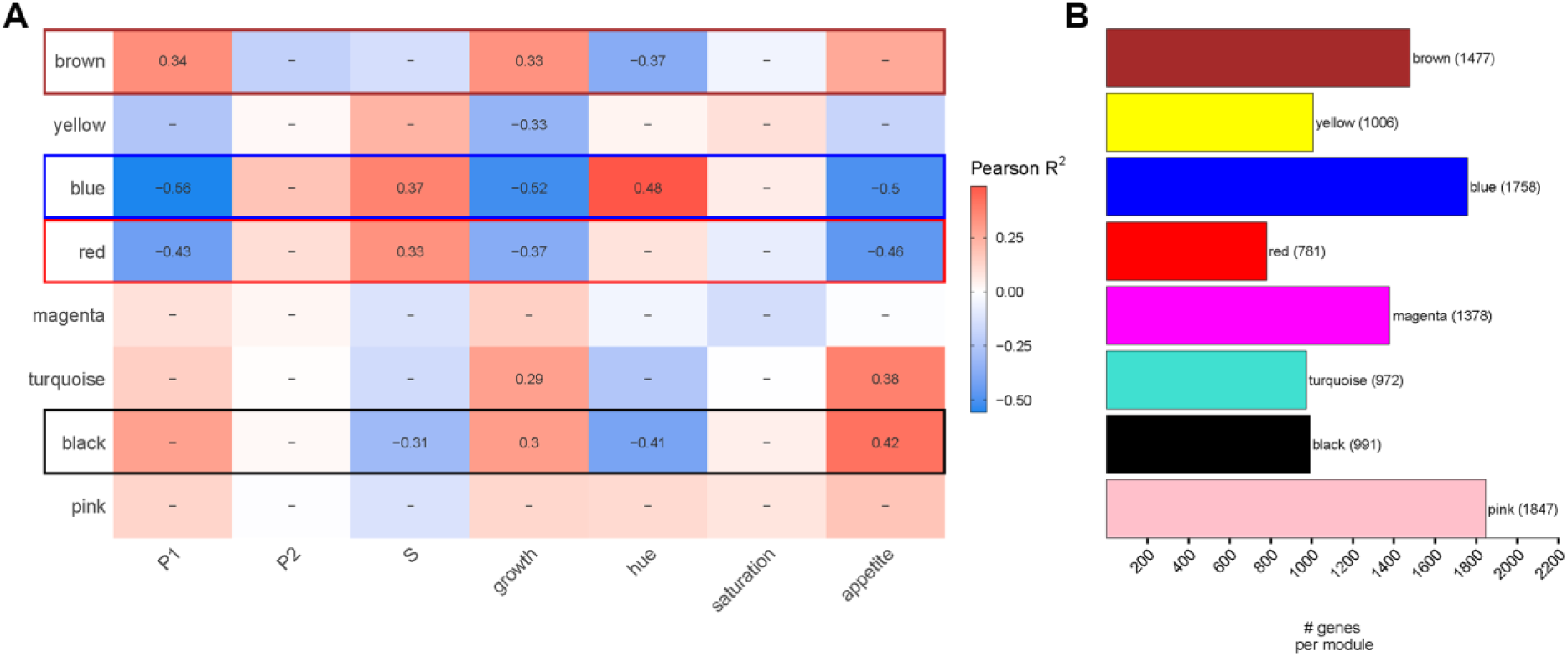
Modules of genes associated with phenotypes of interest. **A)** Heatmap (merging threshold of 0) of Weighted Gene Correlation Network Analysis (WGCNA) to correlate gene expression pattern with social position (P1, P2, S), growth, appetite, orange hue and saturation of individuals. Only significant correlations are shown while non-significant correlations are marked with dash. **B)** The number of genes associated with each module found in WGCNA.

**Supplementary Figure S7:**
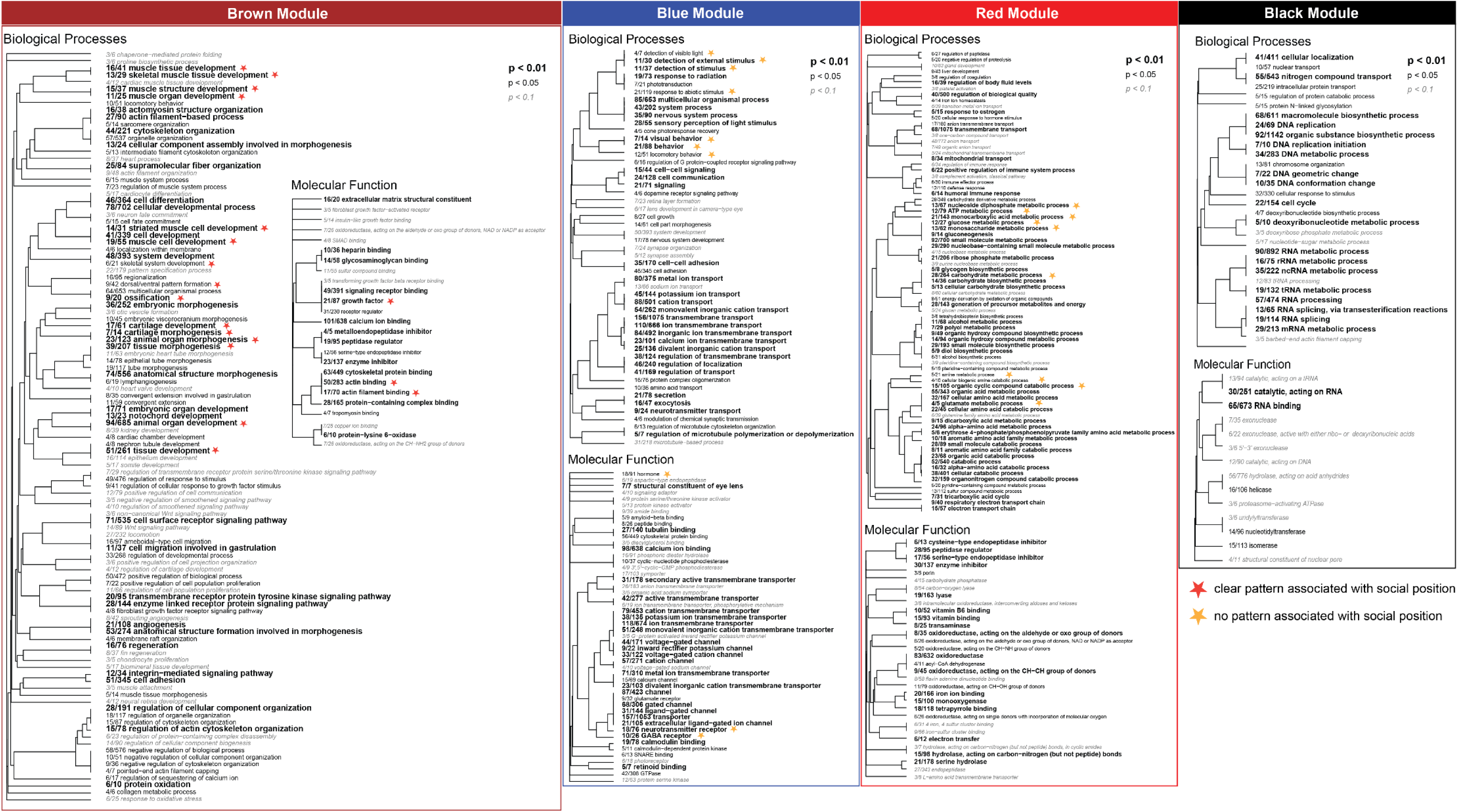
Gene Ontology (GO) terms identified in WGCNA modules showing correlation to phenotypes of interest. Trees generated from GO enrichment analyses using Fisher’s Exact Tests (presence/absence in a module) to identify over-represented pathways within each module across GO divisions of Biological Processes (BP), Molecular Functions (MF), and Cellular Components (CC). Key GO terms, marked with stars, were explored in greater detail by extracting their associated genes and visualizing expression patterns of genes (FDR < 0.05) using heatmaps. Terms with red stars indicated GO terms with clear pattern (heatmap shown in main text), while yellow stars indicate terms which did not show clear patterns of DEGs dependent on social position (heatmaps not shown).

## Growth related genes are upregulated in dominant individuals and not related to size ratio

We have identified genes associated with growth and ossification that showed strong positive correlations with body size. For this set of genes, we tested the hypothesis that the final size ratio of pairs (P2 SL / P1 SL), which is indicative of the amount of remaining conflict (Wong et al., 2007), would predict variation observed in gene expression within social position (Fig. 5B). Visual inspection of the size ratio annotation in the heatmap of these genes (Supplementary Fig. S8), together with a PCA of paired individuals (P1 and P2) based on the same row Z-score values, was used to assess whether remaining conflict explained expression variation (PERMANOVA: p = 0.858; Supplementary Fig. S10A). These results suggest that size ratio between pairs was not associated with gene expression variation within social position.

**Supplementary Figure S8:**
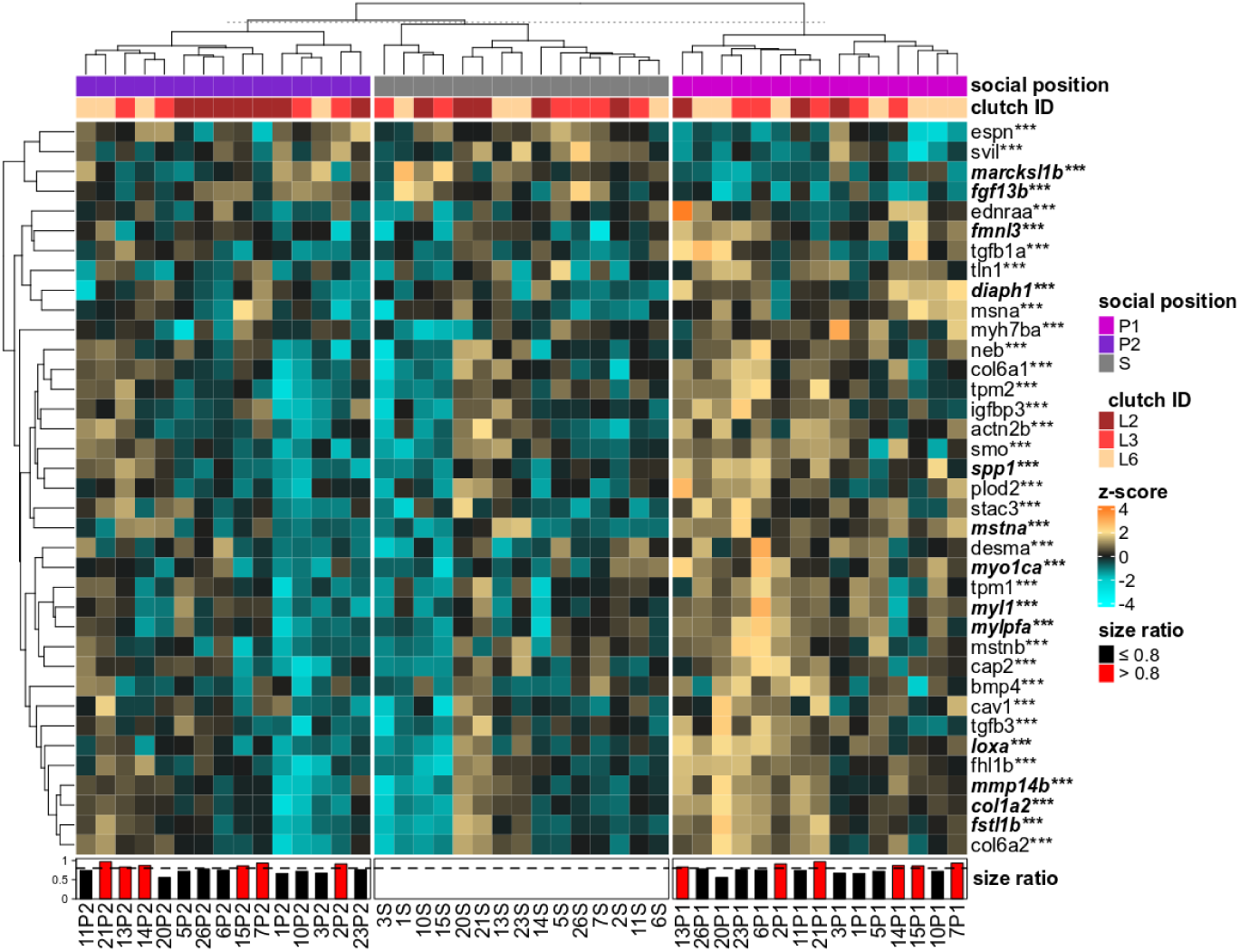
P1 individuals showed upregulation of genes associated with growth and ossification. Heatmap of significant DEGs (FDR p-value ≤ 0.05) of growth and ossification genes in the brown Biological Process (BP) and Molecular Function (MF) GO terms across the three social positions. The bottom panel shows the size ratio (P2 SL/P1 SL) at week 5 for each replicate pair. Black bars indicate pairs with a size ratio ≤ 0.8 (more dissimilar in size), and red bars indicate pairs with a size ratio > 0.8 (more similar in size). The dashed line denotes a size ratio of 0.8. Samples (columns) are grouped *a priori* by social position (dominant, P1; subordinate, P2; solitary, S) and ordered within each group by the column dendrogram, which reflects expression similarity among samples within each social position. The gene (rows) dendrogram reflects relatedness among genes based on expression profile similarity across all samples. Genes that are significantly differentially expressed between individuals of P1 vs P2 are denoted with an asterisk (*), those between P1 vs S with a double asterisk (**), and three asterisk (***) represents both comparisons. Genes shown in ***bold and italics*** indicate significance at an FDR adjusted p-value < 0.1.

## Dominant social role is associated with suppression of satiety signals and altered metabolic pathway activity and not related to size ratio

We have identified genes associated with appetite regulation and metabolism, which were significantly downregulated in P1 individuals compared to P2 and S fish. As above, we tested the hypothesis that the final size ratio of pairs (P2 SL / P1 SL), which is indicative of the amount of remaining conflict (Wong et al. 2016), would predict variation observed in gene expression within social position (Fig. 6). We found no significant effect (PERMANOVA: p = 0.844; Supplementary Fig. S9 and S10B), indicating that size ratio does not explain gene expression differences within social positions.

**Supplementary Figure S9:**
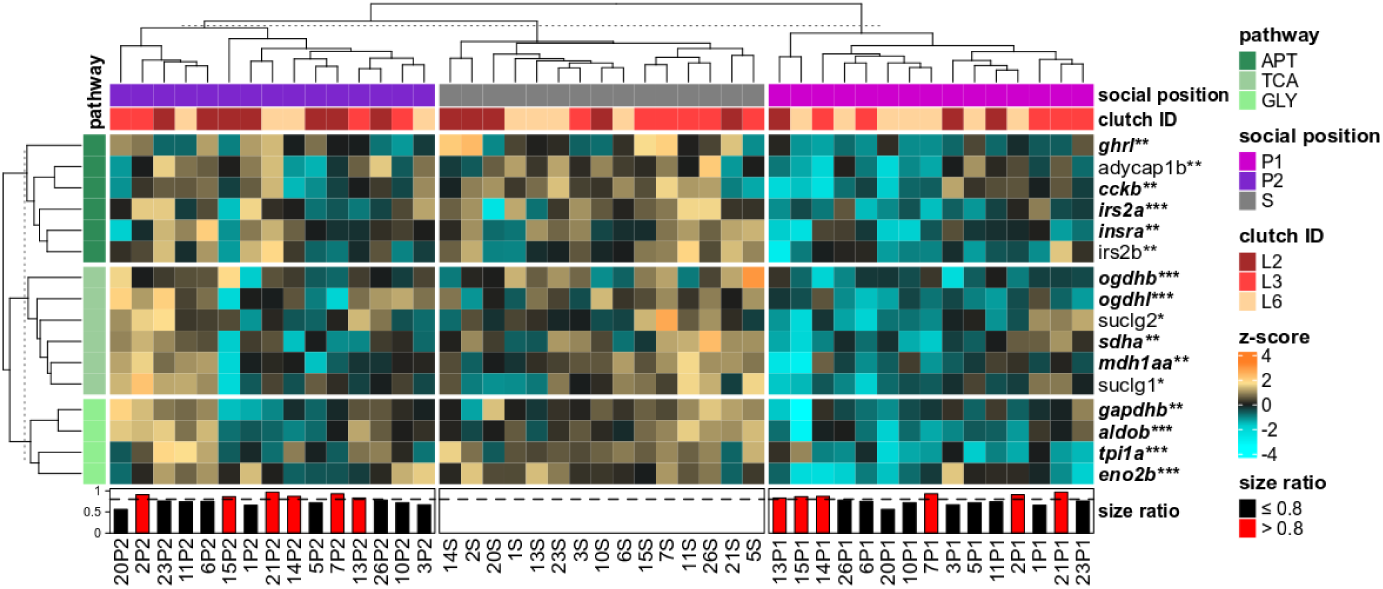
P1 individuals showed downregulation of appetite suppression and metabolic genes. Heatmap of significantly differentially expressed genes (DEGs) (FDR p-value ≤ 0.01) of genes associated with glycolysis (GLY), Krebs cycle (TCA), and appetite regulation (APT), TCA cycle and glycolysis pathways across the three social positions. The bottom panel shows the size ratio at week 5 (P2 SL/P1 SL) for each replicate pair. Black bars indicate pairs with a size ratio ≤ 0.8 (greater size divergence), and red bars indicate pairs with a size ratio > 0.8 (more similar in size). The dashed line denotes a size ratio of 0.8. Samples (columns) are grouped a priori by social position (dominant, P1; subordinate, P2; solitary, S) and ordered within each group by the column dendrogram, which reflects expression similarity among samples within each social position. Genes (rows) are grouped *a priori* by pathway (GLY, TCA, or APT), and within each pathway group, the row dendrogram reflects relatedness among genes based on expression profile similarity across all samples. Genes that are significantly differentially expressed between individuals of P1 vs P2 are denoted with an asterisk (*), those between P1 vs S with a double asterisk (**), and three asterisks (***) represent both comparisons. Genes shown in ***bold and italics*** indicate significance at an FDR adjusted p-value < 0.1.

**Supplementary Figure S10:**
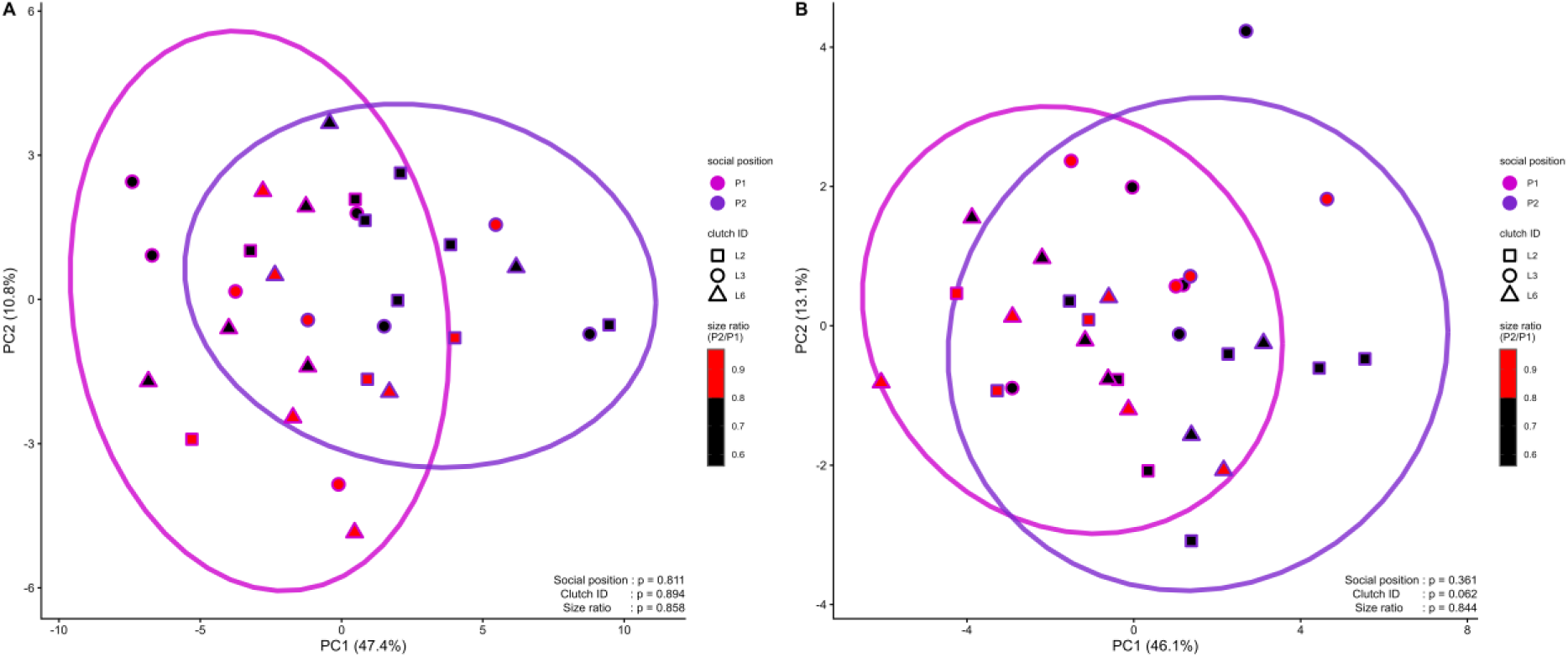
Principal component analysis (PCA) of gene expression in paired individuals (P1, P2) based on heatmaps of A) growth, B) appetite, and metabolic genes. PCA was performed on specific gene sets (growth: Fig 5B; appetite and metabolism: Fig. 6), (P1 and P2; n = 30) using row Z-score scaled expression values, consistent with the heatmap scaling. Each point represents one individual. Ellipses represent 95% confidence intervals around each social position group. A PERMANOVA (*adonis2*; 999 restricted permutations; shown bottom right corners), found that size ratio was not a significant predictor of gene expression variation after accounting for social position and genetic background (A: p_size_ratio_ = 0.858; B: p_size_ratio_ = 0.844).

## Color Data Analysis

We tested the hypothesis that orange hue and saturation will vary depending on the social position of individuals (P1, P2, S), reflecting differences in signaling among individuals (Araújo-Silva et al., 2023). We predicted that dominant and solitary individuals to develop deeper, redder orange hues and higher saturation more rapidly than subordinates.

## Methods

To measure the hue and saturation of orange color of juveniles, fish were photographed in a Petri dish against a white background under a dissecting microscope with a Cannon DS12 camera. Using the white background, images were color-corrected in Adobe Photoshop and color intensity was quantified using RGB values in MATLAB, following a previously established protocol (Winters et al., 2009). Using the macro “AnalyzeIntensity” (Winters et al., 2009), RGB values were obtained for 10 randomly selected points (5 dorsal and 5 ventral) of the orange color area of the second half of the body (from line between soft dorsal rays and anal fins until the line of the caudal peduncle) of all juvenile fish. Using these 10 points, average RGB values were obtained from which hue and saturation were calculated for all individuals. Hue values are indicative of the orange color on the color spectrum, while saturation is indicative of the amount/strength of that given color.

## Results

To test the prediction that hue and saturation will vary dependent on the social position of individuals (P1, P2, and S), a GLMM was run. The best-fit model of hue showed that time (DF = 1, F-value = 639.88, p < 0.001) and social position (DF = 2, F-value = 12.14, p < 0.001) were significant predictors of hue. Post hoc pairwise comparison revealed that both at the beginning of the experiment (t1) and at the end of the experiment (t5), P1 had a significantly different hue compared to P2 and S (both p < 0.05), however, P2 and S showed no significant difference (p > 0.05). Furthermore, post hoc comparison revealed that all fish change in hue, becoming more reddish-orange, over the five-week period (all p<0.001) (Supplementary Fig. S11A). The model, including the fixed effects (social position and time) and random effect (replicate ID), explained 81% of the variation in hue (R^2^c = .811, R^2^m = .779; Supplementary Fig. S11A). The best-fit model of orange color saturation showed that

**Supplementary Figure S11:**
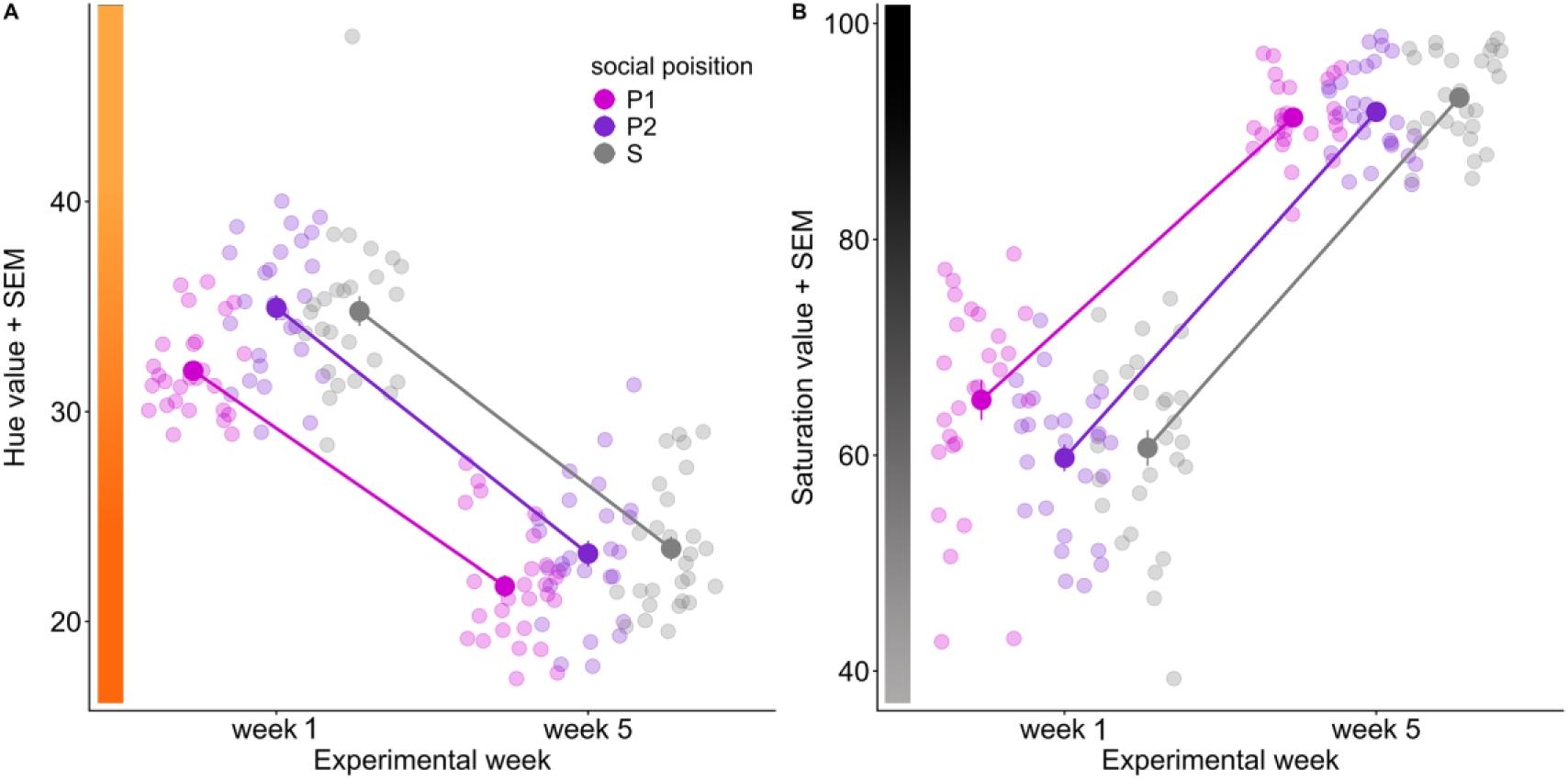
**A)** Hue of individuals of three social position (pair rank 1 = P1; pair rank 2 = P2; solitary = S) over time with mean and standard error of mean (SEM). **B)** saturation of individuals of social position (pair rank 1 = P1; pair rank 2 = P2; solitary = S) over time with mean and standard error of mean (SEM). Data points shown staggered for visual aid, but collected at the same time. time was a significant predictor of saturation (DF=1, F-value = 870.04, p<0.01), all fish showed an increase in saturation over the experiment (Supplementary Fig. S11B).

## Discussion

Here we tested the hypothesis that hue and saturation will vary dependent on the social position of individuals (P1, P2, and S). Our results showed that orange hue value decreases over time as individuals become more reddish orange, a trend that held true irrespective of social position of the individual. Similarly, we found that saturation value of the given color increases over time, irrespective of social position of the individual. We expected to see varying rate of orange hue and saturation development across the three social positions, with individuals taking on dominant roles developing their orange coloration at a different rate from those in subordinate roles. However, we found a mostly uniform change across our individuals. These results do not support the hypothesis that hue and saturation vary depending on social position, contrary to expectations from anecdotal field observations, rather suggest a set of plausible alternative explanations. First, juveniles in our experiment were 4–5 weeks old at the time of introduction to experimental housing, approximately three weeks older than the typical age of wild settlement, raising the possibility that a critical developmental window for color plasticity could have been missed. All fish at the beginning of the experiment had developed moderate orange coloration and thus were already on a color developmental trajectory. Second, all fish experienced a uniform shift in diet and lighting conditions upon entering the experimental tanks, which may have induced consistent changes in coloration across individuals, masking rank-related differences. Indeed, it has been shown before that tank background coloration in combination with varied lighting condition affects development of *A. ocellaris* coloration (Yasir & Qin, 2009). Furthermore, dietary shifts have also been shown to drive changes in coloration, with hue particularly being sensitive to composition of diet (Ho et al., 2014; Tanaka et al., 2016). Moreover, changes in pigmentation may unfold over a longer timeframe than the five weeks monitored. Third, within pairs, subordinate individuals are ultimately expected to assume the rank 2 breeder role and therefore may not face strong pressure to alter coloration, especially if color traits are more relevant for interactions at lower social ranks. Together, these considerations suggest that timing, environmental uniformity, and future reproductive potential may limit the expression or detection of socially mediated color plasticity under laboratory conditions. Future studies should carefully consider experimental design to minimize confounding factors and robustly test the effects of the adoption of social roles on coloration and the effects of coloration on the adoption of social roles.

